# Coupled social-land dynamics and the future of sustainable consumption

**DOI:** 10.1101/2020.09.18.304113

**Authors:** Saptarshi Pal, Chris T. Bauch, Madhur Anand

## Abstract

Dietary patterns have long been a driver of global land use. Increasingly, they also respond to it, in part because of social forces that support adoption of sustainable diets. Here we develop a coupled social-land use dynamics model parameterised for 164 countries. We project global land use under 20 scenarios for future population, income, and agricultural yield. When future yields are low and/or population size is high, coupled social-land feedbacks can reduce the peak global land use by up to 2 billion hectares, if socio-economic barriers to adopting a sustainable diet are sufficiently low. In contrast, when population growth is low or yield is high, reductions in income elasticity can increase peak land use by 100 million hectares. The model also exhibits a regime of synergistic effects whereby simultaneous changes to multiple social and economic parameters are required to change land use projections. This research demonstrates the value of including coupled social-land feedbacks in land use projections.

## Main

From 1961 to 2013 global food demand went up threefold, from 6.4 trillion to 19.4 trillion kilocalories (kcals) per day. This massive increase is attributed to an increase in the world population from 3 to 7.1 billion and an increase in average per capita consumption of food from 1800 kcals/day to 2600 kcals/day over this period [1]. Land is the primary global food supply. In 2013, an estimated land equivalent of 3.5 billion hectares was consumed (72% of agricultural land in that year) while approximately 1.4 billion hectares of land was spent on food wastage [2]. Future expansion in global agricultural land and/or increased intensity of existing farmland usage is therefore a highly probable pathway to meet the enhanced demands of the 21^*st*^ century. However, agricultural expansion and intensification represent major ecological threats, ranging from clearing of forests and habitat fragmentation [3, 4] to increased greenhouse gas emissions [5, 6].

Agricultural intensification faces an uncertain future. From 1961 to 2013, production gains were mostly due to the steady growth in land productivity [5, 1]. Some studies suggest that certain major crops are approaching their yield ceilings in rich countries [7, 8, 9]. There has been a deceleration in yield growth across the globe primarily due to decreasing investment in agricultural research and reduced food production prices in both higher and lower income countries [10]. Slowing intensification may trigger agricultural land expansion to catch up with rapidly growing demand for food.

Mathematical models of sustainable food systems are becoming an increasing topic of research [11, 12, 13, 14, 15, 16, 17]. Research on sustainable pathways for agricultural technologies tend to focus on the supply side of the problem. On the demand side, models often stipulate future demand trajectories that are independent of how the model variables evolve. For instance, sophisticated land system ensemble models that are used to project land use in Intergovernmental Panel on Climate Change (IPCC) reports since their models use scenarios for homogenized dietary consumption patterns as inputs and, as such do not study the dynamics of system-induced drivers of human consumption behaviours [18, 19, 20, 21]. The importance of incentivizing sustainable consumption has been noted [22]. Dietary patterns can heavily influence trajectories of global land use [20, 23, 24, 25, 26] and individuals include environmental factors while making dietary decisions [27, 28] and therefore land use dynamics and socially-influenced dietary choices are coupled to one another through two-way feedback. However, there has been limited investigation into understanding how these shifts in dietary patterns evolve within populations due to social and economic factors, and in particular how they respond to changing land use.

Sustainable consumption is an economically and socially induced process that evolves endogenously in a population and hence can benefit from systematic study using theoretical models. From the individual perspective, adopting a land-sparing sustainable diet may involve paying a cost of losing the personal satisfaction of consuming meat [29, 30]. However, everyone benefits from an individual’s choice to adopt a sustainable diet, since scarce global land use is reduced as a result of that choice. Hence dietary choices represent a public goods game, where individuals may choose to contribute to a common benefit that all members of the group receive, even if they did not make a contribution [31, 32]. Modeling social behavior in public goods games often uses models of social learning dynamics from evolutionary game theory, which captures how individuals learn behaviours from one another [33, 34, 35]. Interest has grown in coupling dynamic social learning models to models of natural processes such as the global climate system [36, 37] and terrestrial ecosystems [38] although social learning dynamic models have not been applied to study coupled dynamics of global land use and dietary decision-making in human populations, to our knowledge.

Here, we introduce a social learning modelling framework for coupling the country-level dynamics of sustainable dietary decision-making under social learning dynamics to country-level land use dynamics. Our objectives are to: (1) show how models of social dynamics and land use dynamics can be coupled to generate novel predictions that are not possible using approaches that treat these systems in isolation from one another, and (2) gain insight into how potential coupled social-land use processes alter both projected global land use and projected dietary trends. Our objective was not to generate projections for policy use. Hence, we opted for a minimal model that was easier to fit to data and gain insight from.

## Model Overview

Our mathematical model describes a social learning process by which individuals learn dietary behaviour from others. Our model captures the two-way feedback between land use and dietary practice: as dietary practices impact global land use, the resulting trends in global land use can, in turn, stimulate behaviour toward more sustainable diets in a closed feedback loop, albeit modified by socio-economic drivers. Details of the model appear in Methods.

For every country, *i*, we define bounds for maximum and minimum per capita land use in year *t* (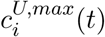 and 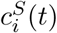 respectively). We classify individuals as having either sustainable or unsustainable diets. Individuals with a sustainable diet consume 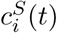 hectares per capita in year *t*. Those with unsustainable diets increase their consumption based on per capita income up to a maximum 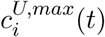. We define *h*_*i*_ as the elasticity of food consumption with respect to income in country *i* (or just, income elasticity of food consumption in *i*). The higher *h* is, the more rapidly consumption changes with income for those practicing an unsustainable diet (see Methods). The estimate procedure for *c*^*U,max*^ and *c*^*S*^ appears in Methods. Beyond 2013 (the last available year in the FAO food balance sheets), these bounds are extrapolated under different scenarios defined by a parameter *f* (a number between 0 and 1). Low values of *f* represent scenarios where future global yields are higher. High values of *f* represent inferior (low) yield futures (see Methods for a mathematical representation of the scenarios).

We assume every country *i* is characterized by a barrier to adopting a sustainable diet, *σ*_*i*_, such that when global land use *L < σ*_*i*_, the perceived costs of a sustainable diet push the population toward an unsustainable diet, while when *L > σ*_*i*_, the population moves toward the sustainable diet. *σ*_*i*_ represents a barrier to achieving population-wide adoption of a sustainable diet due to the combined effects of various psychological, social and economic factors. The rate of dietary change is dictated by *κ*_*i*_, which describes how fast social learning occurs in country *i. κ*_*i*_ is a control knob that determines how often an individuals samples other individuals in the population regarding their diet. If an individual on a non-sustainable diet samples an individual on a sustainable diet and if *L > σ*_*i*_, they switch to a sustainable diet with a probability proportional to the difference *L* − *σ*_*i*_. A similar process occurs for the switch from sustainable to unsustainable diets (see Methods). When *L > σ*_*i*_, the proportion *x* of individuals on a sustainable diet increases as individuals switch from an unsustainable diet to a sustainable diet. The opposite happens in the unsustainable regime. A high value of *κ*_*i*_ can accelerate change in either direction depending on the difference between *L* and *σ*_*i*_.

We use a previously published model [2] to generate country-level land use data based on dietary patterns from 1961 to 2013. We fit our model to these data to estimate *κ*_*i*_, *σ*_*i*_ and *h*_*i*_ for 166 currently existing countries (see Appendix SI Section 1 for methods of parameter estimation and Appendix SI Section 4 for countries included). These estimated parameters were taken as our baseline parameter values. Under the umbrella term ‘agricultural land use’ we included land used for agriculture, pasture and feed generation. Our land calculations excluded land equivalent of food wastage: we accounted only for the land that is used to generate the food that ends up being consumed by the population (See Methods for details). The model parameters, *κ, h* and *f*, are real numbers in the interval (0, 1).

## Results

We make global land use projections for 20 scenario combinations for the 164 countries we analyzed (see Appendix SI Section 4 for details on countries used). For country-level population and income projections, we use the five shared socio-economic pathway (SSP) scenario markers, SSP1 to SSP5 [39, 40]. Each SSP scenario represents a unique storyline for the future that dictates the trajectory of population and income in countries (among other things). Although these scenarios have unique storylines for yield growth, we also show results for different possible future yield trajectories under each SSP scenario. SSP1 is characterized by relatively high income and small population. In SSP2, current trends of population and income continue, and moderate progress is made by achieving income convergence between countries. SSP3–also called the road to regional rivalry–is characterized by an overall high population growth and low income levels in developing countries. The SSP4 future sees high disparity in economic growth rates between high income and low income countries; global growth is less rapid compared to SSP1. In a SSP5 world, economic development is of utmost priority, income growth is high, on average, and it is coupled with strong improvement in education that leads to reduced fertility and hence a relatively small but well-educated population. See Appendix SI Figure 11 for population and income projections under the five SSP scenarios until 2100. For each of the five SSP scenarios, we also explored four scenarios for future agricultural yield: *f* = 0.2, 0.4, 0.6, 0.8, producing a total of 20 scenarios.

### Dynamic social-land feedbacks can partially counteract policy

At the global level, the model shows how social dynamics partially counteract land use impacts caused by other trends such as changing per capita income and population size. The model predicts a net decrease in the proportion of individuals practicing a sustainable diet (*x*, or, ‘sustainable consumers’ hereafter) from 2013 to 2100 in all scenarios, on account of a high average barrier to adopting a sustainable diet (*σ*_*i*_) and increasing per capita incomes (Figure 1b, and see Appendix SI Figure 6 for global distribution of baseline *σ* values). A more rapid decline occurs under SSP5 and SSP1, on account of lower population sizes and thus lower land use in those scenarios creating a reduced perception of need to switch to a sustainable diet (Figure 1a). There are more sustainable consumers higher under SSP3, on account of higher land use in that scenario. In SSP3, due to reduced global income, unsustainable practitioners cannot consume as much as they could have with a higher income. However, this does not help reduce global land use because population size grows fastest under this scenario. Unsustainable practitioners therefore switch to sustainable diets faster because growing global land use exceeds the barrier to adopting a sustainable diet. Their behavioural change is, however, of little avail. Since their unsustainable consumption is not substantially higher than the sustainable level (due to reduced income in SSP3), the effects of this behavioural change are outweighed by high population growth. On the contrary, in SSP1 and SSP5, higher incomes allow higher consumption for the unsustainable practitioners. But low population growth prevents higher per capita consumption from causing a large rise in global land use. As a result, the temporal evolution to sustainable diets is slower in these scenarios.

**Figure 1:**
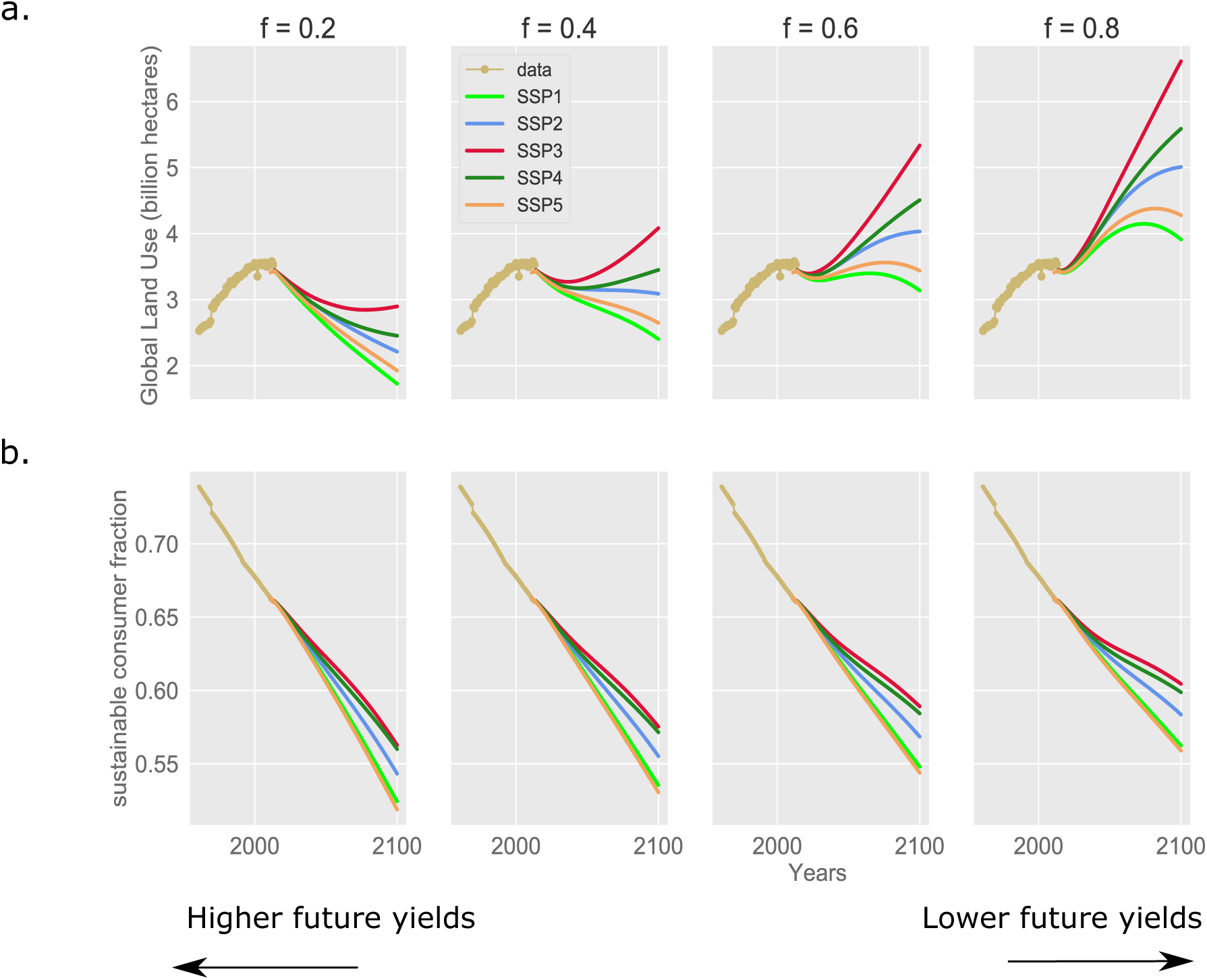
Global land use projections to 2100 under multiple yield and SSP scenarios (a) Global agricultural land use projections till 2100 (excluding land equivalent of food wastage) under 20 scenario combinations. The four columns cover the yield scenarios of *f* = 0.2, 0.4, 0.6 and 0.8. Yellow dots show the time series data for land use from 1961 to 2013. Data from 1961 to 2013 is generated using Ref. [2]. Projections in solid lines begin from 2011 and continue till 2100. **(b)** Model projections of fraction of global population consuming sustainably. See Methods for model definition of sustainable consumption. Yellow dots show time series data for fraction of people consuming sustainably between 1961 and 2013 estimated as in Ref. [2].

Under scenarios of higher future yield (*f* = 0.2 and *f* = 0.4), global land use declines from its 2013 values across most SSPs. The only exception is SSP3 where land use starts to increase again after a period of decline. This occurs because the global population continues growing throughout the 21^*st*^ century under SSP3. Eventually, the effect of population size outweighs the effect of saturating gains in yield. Under scenarios of lower future yield (*f* = 0.6, 0.8), future land use deviates significantly across the SSPs but generally tends upward. We project land use to go as high as 6 billion hectares in the most extreme scenario (*f* = 0.8, SSP3). SSP scenarios with large initial population growth rate (SSP3 and SSP4) do not reach peak land use by 2100. On account of rapidly expanding land use, the sustainable consumers decline less rapidly than in the higher future yield scenarios, but the overall trend is still downward.

Taken together, these results show how changes in parameters such as population size and per capita income can cause a social response that partially counteracts those changes. For instance, SSP5, despite being the most sustainable scenario in other respects, does not exhibit the strongest transition to sustainable diets because the reduced population size in that scenario causes a reduction in land use required, and thus reduces the perceived need to transition to a sustainable diet. Similarly, higher agricultural yields reduce land pressure, and thereby also reduce the perceived need to transition to a sustainable diet. Scenario combinations involving higher future yield and/or SSPs with lower population size cause sustainable consumers to decline, which means that land use ends up being higher than it would be without this feedback between land use and dietary choices.

### Continental and country-level land use projections

Projections broken down by geopolitical region reveal significant heterogeneity behind the global trends (Figure 2). Europe and Oceania exhibit an increase in sustainable consumers and a decrease in land use across all SSPs. This is because countries in Europe and Oceania have lower inferred barriers to adopting a sustainable diet (*σ*_*i*_) compared to the rest of the world (Appendix SI Figure 6). With respect to evolution of global land use, they always remain in the regime where sustainability is the dominant behaviour with higher utility. The relative ordering of land use by SSP we saw in the global projections remains consistent at the continent level. These projections also show that an increase in sustainable consumers will not necessarily lead to a decrease in land use, even if that is the general trend.

**Figure 2:**
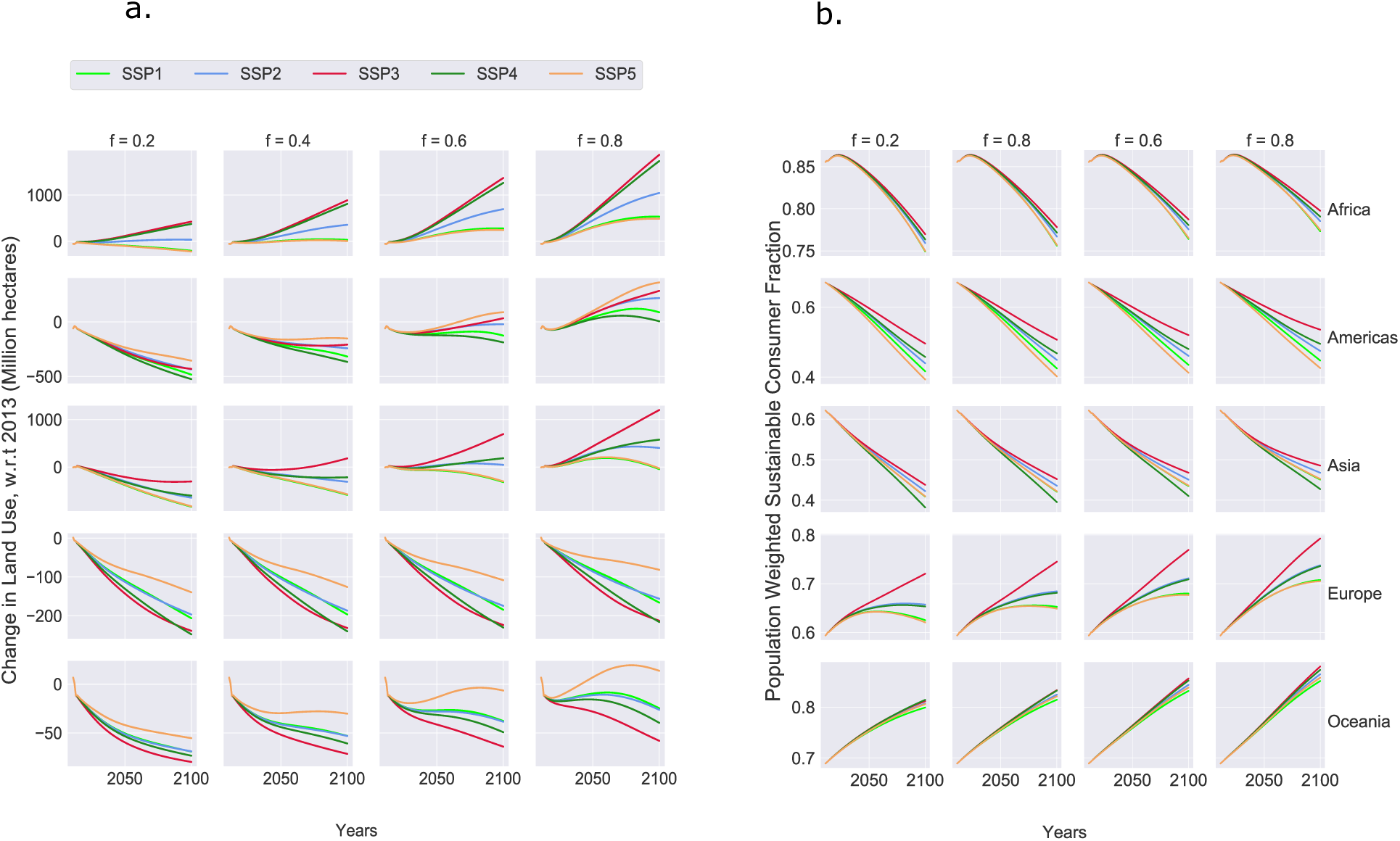
Global land use projections to 2100, broken down continent-wise, reveal heterogeneity under sustainable behaviour in baseline conditions. **(a)** Projections of change in agricultural land use with respect to 2013 (excluding land equivalent of food wastage) broken down continent-wise into five major regions - Africa, Asia, Americas, Europe and Oceania (see Supplementary Section 5.2 for division methodology). Projections are shown for 20 scenario combinations (combinations of 5 SSP scenarios and 4 *f* scenarios). **(b)** Model projections for fraction of regional population consuming sustainably for 20 scenario combinations. See Methods and Supplementary for formal definitions of sustainable consumption. Only Europe and Oceania show a rise in sustainable consumers over the projecting period (2011 - 2100).

For example, in certain scenarios, Africa, Asia and the Americas show a decrease in land use (with respect to 2013) while the fraction of sustainable consumers also declines, on account of growth in agricultural yield outweighing the effects of income and population growth. For these regions, projections under the SSP3 scenario shows the highest use of land. This is because, for them, population projection under SSP3 is the highest among all SSP scenarios (unlike Europe and Oceania where it is the lowest). That, coupled with a steady decline in sustainable consumers in their population, results in the fastest change in land use. For these regions, the average socio-economic barrier to adopting a sustainable diet is always higher compared to the evolution of global land use in all of the 20 scenario combinations. This indicates that future yield, income and population do not drive the growth of sustainable consumers and the decline of land use identically. The feedback loop between land use and dietary behaviours in our country-level model gets scaled up to the regional level, too. In other words, each region shows a unique behavioural response to change in global land use because of its unique social setting.

Some countries with relatively small population sizes are projected to emerge as front runners of global land use (Figure 3). The correlation between population and land use is not absolute, however (Figure 3b). In 2013, the countries that had a comparatively lower population but high land use were Kazakhstan (population, 18 million), Saudi Arabia (33 million), Australia (25 million) and Argentina (45 million). Ranking projection of land use shows that it is likely the fourth spot, currently occupied by Russia, will be taken over by Saudi Arabia by 2050 (which, in 2013, occupied the fifth spot in land use) (Figure 3a). In 2013, the two countries consumed comparable areas of global agricultural land for their respective demands (123 million hectares for Russia and 105 million hectares for Saudi Arabia). The primary reason for Saudi Arabia overtaking Russia can be identified from Figure 3c. Russia sees an increase in sustainable consumption over the time horizon, while Saudi Arabia sees a decrease. In all the scenarios, their baseline parameter value of *σ* (barrier to adopting a sustainable diet) places them on opposite regimes of behavior with respect to evolution of global land use.

**Figure 3:**
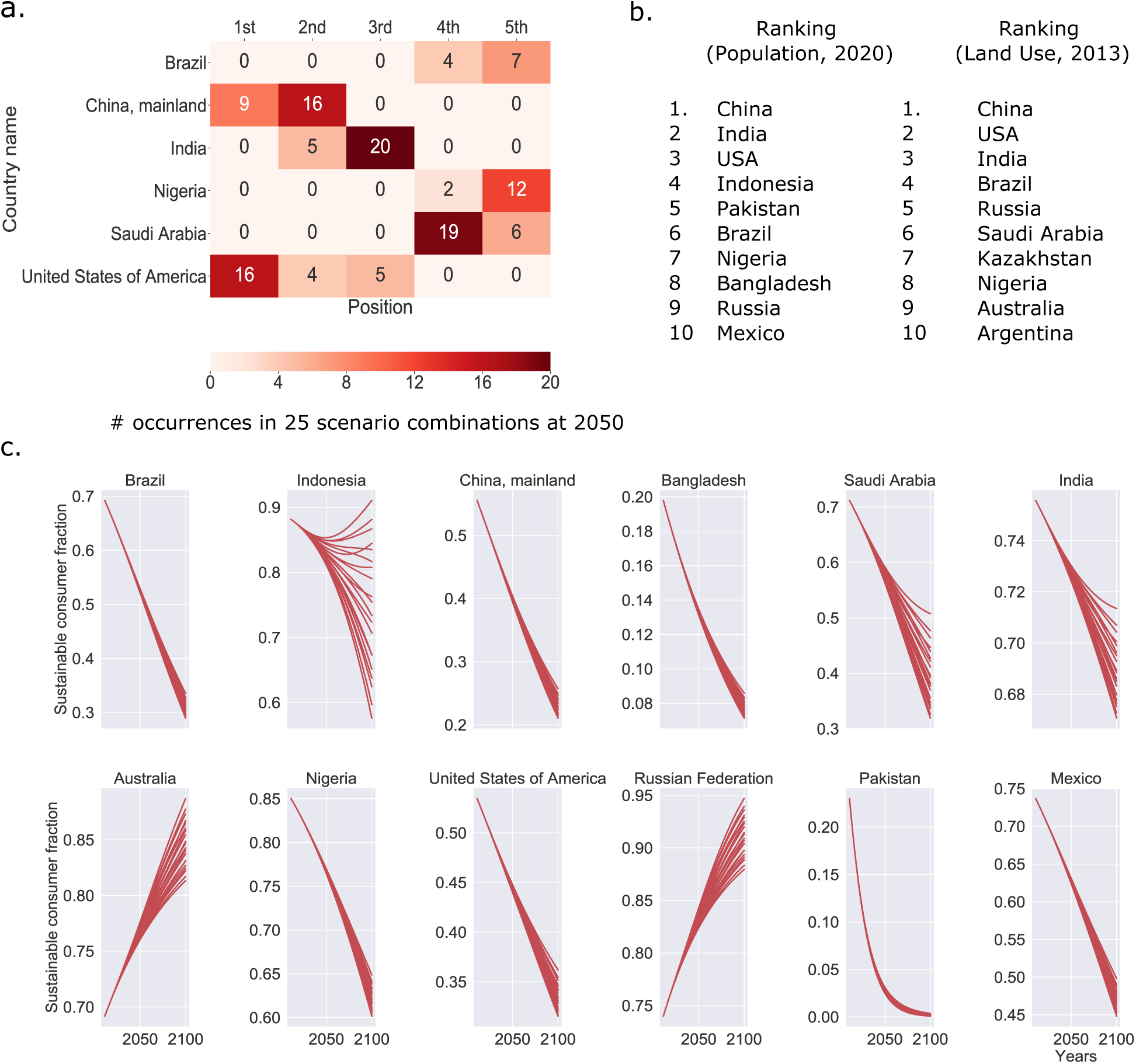
Countries with high population and high land use show general trend of reducing sustainable behaviour under baseline parameters. **(a)** Ranking of countries based on their total land use across 25 scenarios at 2050. Six countries land at least once inside the top 5 positions in 25 scenario combinations (combinations of 5 SSP scenarios and five yield scenarios: *f* = 0.2, 0.4, 0.6, 0.8 and 1). The heat map indicates the number of appearance of a country at a particular ranking position. China and the USA dominate the first two spots while India, Saudi Arabia and Nigeria dominate third, fourth and fifth positions respectively. **(b)** Table showing ranking of countries based on their population (2020, data) and land use (2013, data generated from model in Ref. [2].). **(c)** Model outputs of sustainable consumers for the twelve countries that occupy spots in either of the rankings in b.

### Synergies can reduce peak global land use

We found that socio-economic factors as represented in our model–the social learning rate (*κ*), the barriers to adopting a sustainable diet (*σ*), and income elasticity (*h*)–have very large impacts on peak global land use, often ranging in the giga-hectares (Figure 4). This is particularly true when higher incomes, higher population sizes and lower future yields force individuals to make a choice between sustainable and unsustainable diets in the face of rapidly expanding global land use. In contrast, when land use does not expand as rapidly due to lower population sizes or higher yields, the perceived need to switch to a sustainable diet is less.

**Figure 4:**
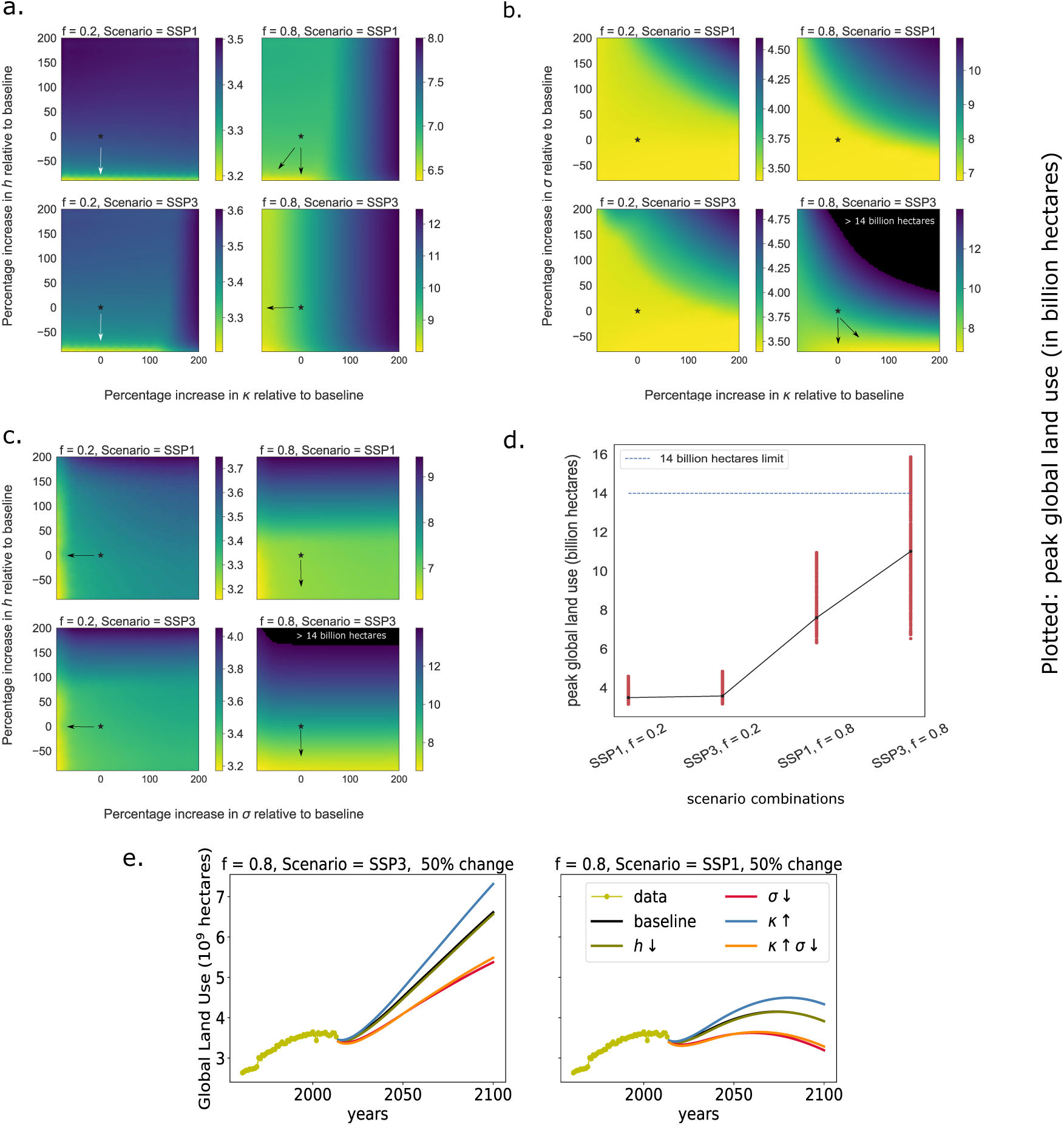
Variations in social parameters from baseline level impact global land use. Model output of peak global land use at four scenario combinations - *f* = 0.2, SSP1, *f* = 0.2, SSP3, *f* = 0.8, SSP1 and *f* = 0.8, SSP3. Model projections are evaluated at parameters deviated from their baseline settings. The black star in each plot indicates the position of the baseline parameter in the heat map. Heat map for peak land use projection with deviations in **(a)** *κ* (social learning rate) and *h* (income elasticity of food consumption), **(b)** *κ* and *σ* (barrier to adopting a sustainable diet), and **(c)** *σ* and *h*. All other parameters held at baseline values. The unit for the color bar in the heat-map is billion hectares. **(d)** Peak global land use values from (a)-(c) plotted versus the scenarios. **(e)** Model time series of global land use in the 21^*st*^ century in scenarios of low global yield; parameters varied within 50% of baseline value.

When future yields are lower (*f* = 0.8), the peak global land use is much more sensitive to social processes than when future yield is higher (*f* = 0.2) (Figure 4). Low yield means rapidly expanding land use, which in turn stimulates a social response in favour of wider adoption of a sustainable diet. Hence in this scenario, changes in social parameters governing the pace and desirability of change have large impacts on land use. When future yields are low, population growth also becomes a determining factor in assessing the effectiveness of varying social parameters (Figure 4d). In contrast, when future yield is high, land use is lower even though more individuals are practicing an unsustainable diet, and thus changes to parameters governing pace and desirability of a sustainable diet have less impact.

In scenarios where future yields are lower, an increase in the social learning rate (*κ*) leads to high peak global land use due to faster conversion to unsustainable consumption as per capita income rises (Figure 4a). Globally, the barrier to adopting a sustainable diet is too high for sustainability to spread in populations even when global land use increases quickly. In the model, if a sustainable consumer samples their population very often (high social learning rate, *κ*), they are easily tempted to shift to unsustainable consumption because they see an increased expected utility in switching. The only way to reduce this effect is to reduce the barrier to adopting a sustainable diet (lowering *σ*, Figure 4b). This could be possible by incentivizing consumption of plant protein by reducing the market price of animal protein substitutes or increasing public knowledge about health and environmental implications of a high meat diet. Once the barrier to adopting a sustainable diet is sufficiently low, social learning rates assist in lowering the peak global land use (Figure 4b). When *σ* is lowered, sustainable consumption becomes the dominant behaviour due to its higher utility. In this case, a considerable amount of land is saved even in scenarios where global population growth rate is high (SSP3) and future yields are inferior (*f* = 0.8).

If global land use evolves very slowly due to slow population growth (SSP1, SSP5) and high global yield (*f* = 0.2), the model predicts that sustainable consumption never becomes the dominant behaviour at the global level. This is because *L* always remains significantly lower to the baseline values of *σ* in these scenarios. Even with sizeable changes in social parameters, *κ* and *σ*, only an insignificant increase in sustainable consumers is achieved. As a result, there is no direct impact on peak global land use. As global land use change is small in these scenarios (and sometimes negative, see Figure 1a), there is not enough incentive for individuals to even pay a lowered cost to being sustainable. In such a setting, the key to reducing global land use lies in the consumption patterns of highly prevalent unsustainable consumers. Since global average income is high in these scenarios, an increase in income elasticity can potentially cause negative impacts on global agricultural land use (Figure 4a, 4c).

However, in the least optimistic scenarios (high population growth rate and low future yields), certain variations of social parameters from their baseline values can alter peak global land use by approximately two billion hectares (twice the size of China). Depending upon socio-economic and yield growth scenarios, the optimal strategy for lowering peak land use changes. Although it is always beneficial to reduce the barrier to adopting a sustainable diet, there can be scenarios where better gains are achieved by modulating the consumption patterns of unsustainable consumers. Varying all three social parameters guarantees a synergy in terms of lowering of peak land use in the 21^*st*^ century, irrespective of socio-economic scenario, while varying only two of the three parameters sometimes has little effect.

## Discussion

Individual diets are influenced by complex social factors such as religion, concern for health, urbanization, female participation in labour, food prices, and sustainability practices [41, 42, 43, 44]. Several of these factors imply a two-way feedback between land use and dietary decisions. Here we focused on the effect of ballooning global land use as a stimulus for individuals to adopt more sustainable diets, against a backdrop where rising incomes also permits individuals to opt for unsustainable diets instead by eating more land-intensive foods such as meat. We subsumed other factors in decision-making into our phenomenological parameters at the social (*κ, σ*) and individual (*h*) level that we inferred from data.

We showed how coupled social-land dynamics can have giga-hectare impacts on land use, especially when future yield is low and/or population size is high, and we explored changes to social parameters that minimize future land use under various scenarios for socio-economic development pathways and future agricultural yield. We found that reducing barriers to adopting sustainable diets is an important way to reduce peak global land use. Increasing social learning rates holds the potential to accentuate the mitigating effect of reducing socio-economic barriers (a simultaneous effect shows a reduction of 2 billion hectares in peak global land use). Increasing social learning can result in negative effects if no improvements in lowering barriers are made, however.

Our minimal model made simplifying assumptions that could impact its land use projections. For instance, we did not include aquatic sources of food, we ignored the influence of institutions, and we assumed a binary classification of consumption behaviour. A future extension of our model could include aspects of population heterogeneity such as a continuous behavioural spectrum along with age and gender structure. Future work could also explore the effects of social norms in order to determine how social inertia can accelerate or decelerate behavioural changes, as well as social learning between countries. For the purpose of simplicity in working with country level data, we also assumed homogeneous behaviour within each country by assigning unique parameter values to every country, and this could be relaxed in future research. Similarly, given the enormous greenhouse gas impacts of livestock [45], a future social process model would take into account the perceived risk of climate change in modeling the behavioural drivers of a population.

Future research in coupled social-land use models can incorporate increasing sophistication to deepen our understanding of social processes around dietary choices and land use dynamics, as well as their interaction with other socio-economic factors and other environmental dynamics such as climate change. These models could inform land use projections and deepen our insights into relevant processes, by incorporating the driving mechanisms behind our dietary choices and accounting for how they respond to changes in land use and socio-economic variables.

## Methods

### Coupled social-land Use Model

For a country *i* and year *t*, we assume two possible diet types: sustainable and unsustainable for the entire population. The sustainable diet type requires 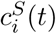 hectares per capita to generate while the unsustainable diet requires 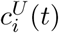 hectares per capita. By definition, 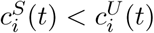 for all *i* and *t*. We make the assumption that the sustainable diet is within reach of anyone in a country irrespective of income whereas unsustainable is aspiration-only. When income is small, individuals aspiring to an unsustainable diet are only able to include occasional land-intensive items in their diet, but as their income rises, they include more. We represent this behaviour with the following equation:

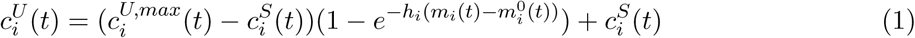

Where 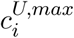 is upper limit of consumption by the unsustainable practitioners, *m*_*i*_(*t*) is the average income of the population and 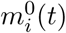 is the minimum income that can afford the sustainable diet at *i* in *t*. The parameter *h* denotes the elasticity in the behaviour of unsustainable practitioners. If *h* is large, *c*^*U*^ grows towards *c*^*U,max*^ faster with income as compared to when *h* is small. Note that when average income *m*_*i*_(*t*) equals 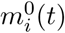, the entire population consumes sustainably; that is, they consume 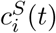 hectares per capita. The per capita consumption of practitioners of unsustainable diet, 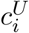, approaches *c*^*U,max*^ asymptotically as the difference between *m*_*i*_ and 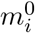 gets higher. Our assumption that meat and dairy consumption increases with income has been explored and identified in earlier papers like [46, 41].

Let *x*_*i*_(*t*) and 1 − *x*_*i*_(*t*) be respectively the proportions of the population that are practitioners and non-practitioners of sustainable diets in *i* at *t*. The average per capita consumption of the population can then be defined as follows:

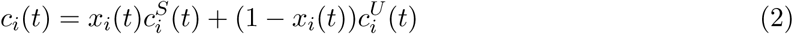

If *P*_*i*_(*t*) is the population of *i* in *t* then the land used due to dietary consumption of population *i* at *t* is *P*_*i*_(*t*)*c*_*i*_(*t*). Global land use, or, the land used due to consumption by the entire population of the globe at *t* can then be defined as the sum of land consumed by all the nations in the world at *t*:

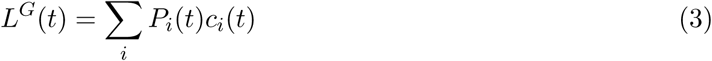

We use imitation dynamics from evolutionary game theory to describe the time evolution of *x*_*i*_. The utility gain for changing from an unsustainable diet to a sustainable diet for the baseline model is given by

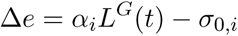

Hence, as the impact function *L*^*G*^(*t*) rises over time due to growing incomes, there is a growing incentive for individuals to switch to a sustainable diet, according to a proportionality constant *α*_*i*_. The rate of switching becomes faster as the difference between *α*_*i*_*L*^*G*^ − *σ*_0,*i*_ grows and vice versa. However, this behaviour to switch to sustainable practice is only effective when *α*_*i*_*L*^*G*^ is greater than *σ*_0,*i*_. When *α*_*i*_*L*^*G*^ is less than this threshold, *σ*_0,*i*_, the proportion of unsustainable practitioners grows, the rate being determined by the absolute difference between *α*_*i*_*L*^*G*^ − *σ*_0,*i*_. We call the parameter *σ*_0,*i*_, the socio-economic barrier to adopting a sustainable diet in *i*. Assuming a social learning rate of *κ*_0,*i*_ for *i* we can write the evolution of sustainable practitioners as follows:

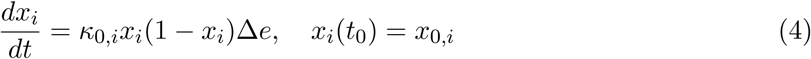

After some rescaling of parameters we obtain:

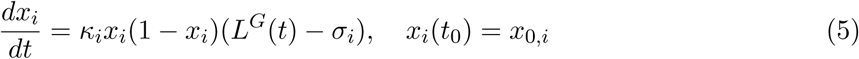

Where *κ*_*i*_ = *κ*_0,*i*_*α*_*i*_ and *σ*_*i*_ = *σ*_0,*i*_*/α*_*i*_ are the rescaled parameters. We refer to the rescaled parameters *κ*_*i*_ and *σ*_*i*_ with their original names. That is, *κ*_*i*_ is social learning rate and *σ*_*i*_ is the barrier to adopting a sustainable diet in *i*. When global land use *L*^*G*^(*t*) exceeds *σ*_*i*_, unsustainable practitioners switch to sustainable behavior at a rate which is determined by *κ*_*i*_, the existing proportion of sustainable practitioners and the absolute difference between global land use and *σ*_*i*_. When global land use is less than *σ*_*i*_, sustainable practitioners switch to unsustainable behaviour through the same mechanism.

### Method for calculating 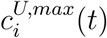 and 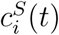

The upper bound of per capita consumption, *c*^*U,max*^, is calculated by assuming that the maximum diet is the one that allows highest intake of items that belong in the meats and dairy diet groups. Similarly, for *c*^*S*^, we assume that sustainable diet is the one that allows least consumption of items in those groups. Our assumption is backed by numerous studies that have found meat intensive diets to be environmentally unfriendly and land-intensive [18, 47]

*c*^*U,max*^ and *c*^*S*^ can be calculated between 1961 and 2013 for countries whose data is reported in FAOSTAT’s food balance sheets [1]. We categorize each of the 21 food items listed in the food balance sheets into one of the seven groups of diet - fruits, vegetables, grains, meats, dairy, oils and sugar.

For every country *i*, we calculate its maximum possible diet by replacing its average consumption of items in the ‘meats’ and ‘dairy’ groups (in kcals/capita/day) with the consumption values of the countries that consumed the most of those items that year. Similarly, for the minimum sustainable diet, we replace them with the consumption values of countries that consumed the least of those items in that year. Values for the remainder of the diet (i.e the other groups - fruits, vegetables, grains, sugar, oils), remain the same as reported data. An example of such a construction is shown in Appendix SI Table 1). The method of evaluating these bounds are explained with more detail in Appendix SI Section 1.1.

Once these hypothetical maximum and sustainable diets are constructed for a country *i*, we use the model developed in Ref. [2] to calculate the total land required to generate that per capita dietary demand for the population of *i* in *t* (see Appendix SI Section 2 for an overview of this model). We divide the output of the model with the population of *i* at that year to obtain per capita land use equivalent of the hypothetical diet (*c*^*U,max*^ if maximum diet, *c*^*S*^ if sustainable diet).

In order to evaluate these values for years beyond 2013 (for purpose of projections), we use an extrapolating parametric function (See Method section for *f* scenarios).

### Definition of land use: Data and Methods

We use the model developed in [2] to generate the country-level time series data of average per capita land use between 1961 and 2013. The model is described briefly in Appendix SI Section 2. The UN FAOSTAT data-set also provides country level data for land used on agriculture and pasture land. However, this is not the same as our definition of ‘land use by *i*’. This is because countries are not entirely self-dependent in providing for their food demand. Consume in *i* can be partly produced in *j* and vice-versa. Since the model in [2] accounts for differential yields of food sources, the data for per capita land use, as generated by model in [2], accounts for land used from across the globe to provide for the consumption in *i*. If two countries have similar dietary consumption, the country which has a lower effective yield has higher value of per capita consumption than the country which has a higher value of effective yield.

In all our projections and analysis, we consider land that is required to generate the food that ends up being consumed by humans. Land equivalent of food wastage is not considered in our calculations. The data reported by UN FAOSTAT’s land statistics division [48] accounts for land used for all agricultural purposes. This includes land equivalent of food wastage. In Appendix SI Figure 1, we see the quantitative difference between their time-series and our global model output. FAOSTAT estimated that 1.4 billion hectares were lost due to food wastage in the year 2007 [49]. This number matches exactly with the difference between the two series at 2007 in Appendix SI 1.

### Population, income and *f* (yield) scenarios

We borrow the SSP scenarios (Shared Socioeconomic Pathways) introduced in [39] for projecting population and income to 2100. A number of existing models are compiled in the SSP Public Database hosted by the International Institute for Applied System Analysis (IIASA). Among them, we choose the OECD Env-Growth Model [40] for obtaining future projected values of country level population and income. In Appendix SI Section 4 we discuss the inclusion procedure of countries in our analysis. There we provide reasons for the exclusion of certain countries from the analysis. The choice for OECD Env-Growth was made because it covers projections for maximum number of countries among the existing models.

The bounds for maximum and minimum per-capita consumption (*c*^*U,max*^ and *c*^*S*^) are projected into the future with a parametric function. The parameter *f*, a number between 0 and 1, represents scenarios of yield future. We now explain the meaning of a yield scenario parameterized by *f*. If the trend of *c*^*U,max*^ and *c*^*S*^ between 1990 to 2013 is decreasing (which is more often than increasing), the series can at least reach *f* times its 2013 value in the future. Similarly, if the trend is increasing, it can reach at most 1 + *f* times its 2013 value in the future. The rate at which a projected curve (either *c*^*U,max*^ or *c*^*S*^) reaches towards its bound is determined by its rate between 1990 and 2013.

Let *c* be the concerned time series that we wish to project till 2100 using our parametric function. The series *c* can either be *c*^*U,max*^ or *c*^*S*^ for a country *i*. The series is always defined between 1961 and 2013. First, we fit an exponential of form *y* = *ae*^*bt*^ to a truncated *c* series. This truncated version of *c* is the time series of *c* from 1990 to 2013. If *b <* 0 we call the series trend decreasing and if *b >* 0 we call the series trend increasing. Here, *a* and *b* are constants. We extrapolate the time series *c* till 2100 (starting from 2013 onward) using the following equations:

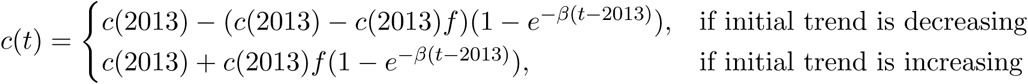

Here *f* is the tune-able parameter - a real number between 0 and 1 that defines the future yield scenario. For the above equation, *t* is always greater than 2013. The exponent *β* is adjusted such that continuity is maintained at 2013 between the initial trend, *ae*^*bx*^, and the projected trend *c*(*t*).

That is,

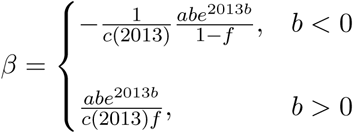

In Appendix SI Figure 9, we show two examples of *c*^*U,max*^ and *c*^*S*^ projection till 2100 using the above method. The two countries that are chosen as examples are USA and Netherlands. USA shows a decreasing initial trend (b *<* 0) whereas Netherlands shows an increasing initial trend (b *>* 0).

If we assume that maximum and sustainable dietary distributions (in kcals/capita/day) for countries remain constant from 2013 onward, *f* scenarios represent scenarios of yield future. Then, a low *f* value represents improvement towards high yield values. A high *f* value represents deceleration of yield rates, causing them to converge to inferior future values.

### Parameter plane analysis

The three social parameters, *κ, σ* and *h* are varied from their baseline values in a pairwise fashion while keeping the third parameter fixed at the baseline setting. Every parameter is varied from -100% to 200% of its baseline value. That is, if *α* is a social parameter, we vary it from 0 to 3*α* while conducting this analysis.

Since we begin projecting at 2011 and continue till 2100, we make the corresponding changes in social parameters at 2011 and keep them that way for the entirety of the projecting period. We make equal percentage changes to social parameters of all countries included in our model. In the parameter planes, we observe the effect of changes in parameter values on peak global land use attained between 2011 and 2100.

We show results for four scenario combinations - i) SSP1, *f* = 0.2, ii) SSP3, *f* = 0.2, iii) SSP1, *f* = 0.8 and iv) SSP3, *f* = 0.8. In all the parameter planes, the colors represent the value of peak global land use (based on an accompanying color-bar). All units of peak global land use are in billion hectares. Baseline parameters are marked by a black star (no change) in each parameter plane. Arrows indicate direction towards least peak global land use.

