## Supplementary Appendix for "Coupled social-land dynamics and the future of sustainable consumption"

#### 1 Method of parameter estimation

For each country $i \in I$, our land-use model is represented by the following coupled equations:

$$c_i(t) = L_i(t)/P_i(t) = x_i(t)c_i^S(t) + (1 - x_i(t))c^U(t) \tag{1}$$

$$c_i^U(t) = (c_i^{U,max}(t) - c_i^S(t))(1 - e^{-h_i(m_i(t) - m_i^0(t))}) + c_i^S(t) \tag{2}$$

$$L^G(t) = \sum_i P_i(t)c_i(t) \tag{3}$$

$$\frac{dx_i}{dt} = \kappa_i x_i(1 - x_i)(L^G(t) - \sigma_i), \quad x_i(t_0^i) = x_{0,i} \tag{4}$$

Note that equations 3 and 4 couple together the model equations for all $i \in I$. The goal of the parameter estimation process is to evaluate the parameters $\kappa_i, \sigma_i, h_i$ and $x_{0,i}$ for a country $i$ given we have the time series data of $i$'s per-capita land use. We define the time period over which we parameterize our model as $[t_0^i, t_f^i]$. For most countries, $t_0^i = 1961$ and $t_f^i = 2013$. There are some exceptions like the Russian Federation for which $t_0^i = 1992$. The yearly data available to us from [1] is within the range of 1961 to 2013. For the parameter estimation part, we **do not** use equation 3 to feed the differential equation in equation 4. Instead, we read $L^G(t)$ from the data. We generate this data from the model in [2]. The model is discussed in detail in section 2 of this document. This time series is shown in Supplementary Figure 1 as the yellow time series.

The parameter estimation process for $i$ also requires as input in the time series $c_i^{U,max}(t)$, $c_i^S(t)$ and $m_i^0(t)$. These time series should cover all the years over which parameterization is being performed. That is, they should be defined in $[t_0^i, t_f^i]$. In the subsections 1.1, 1.2 we discuss methods to obtain these time series. In subsection 1.3, we discuss a method that reduces the number of free parameters from four to three by evaluating $x_{0,i}$ from $h_i$ and the first data point. In subsection 1.4, we discuss the optimization technique we use to parameterize our model. In subsection 1.5, we discuss methods to determine the parameterization period, $[t_0^i, t_f^i]$ for $i$. In section 4, we discuss how the set $I$ is constructed for this work. We explain reasons for inclusion and exclusion of countries from the set $I$.

##### 1.1 Method for generating $c_i^{U,max}(t)$ and $c_i^S(t)$ time series

We define $c_i^{U,max}(t)$ and $c_i^S(t)$ to be the theoretical maximum of per capita land use of non practitioners of sustainable diet and the per capita land use of sustainable practitioners in $i$ at $t$ respectively. Let us represent the Rizvi et al. model as a function, $R(.)$, that maps a diet $\mathbf{D}$, a country $i$, and a year $t$ into a land use value for the country $i$ at $t$. It implies that if the population of $i$ at $t$ consumed the per capita diet $\mathbf{D}$ on average, $R(\mathbf{D}, i, t)$ hectares of land would have been spent, globally, to generate the demand. We can then define $c_i^{U,max}(t)$ and $c_i^S(t)$ accordingly:

$$c_i^{U,max}(t) = R(\mathbf{D}^{max,i}(t), i, t)/P_i(t)$$

$$c_i^S(t) = R(\mathbf{D}^{S,i}(t), i, t)/P_i(t)$$

| Item | Food Group | Highest (kcals/capita/day) | Lowest (kcals/capita/day) |
|---|---|---|---|
| Bovine Meat | Meats | Argentina (342) | Liberia (2) |
| Sheep and Goat Meat | Meats | Mongolia (327) | Congo (1) |
| Pig Meat | Meats | Hong Kong (385) | Guinea (1) |
| Poultry Meat | Meats | St. Lucia (273) | Chad (2) |
| Eggs | Meats | Japan (76) | Cameroon (1) |
| Milk | Dairy | Iceland (562) | Liberia (5) |
| Butter (Ghee) | Dairy | New Zealand (187) | Angola (1) |

Table 1: Highest and Lowest Meats and Dairy item consuming nations in 2010 and their respective average per capita caloric consumption from UN FAOSTAT Food Balance Sheet [1]

Where $\mathbf{D}^{max,i}(t)$ is a hypothetically large diet that is considered as the theoretical maximum diet for non-practitioners and $\mathbf{D}^{S,i}(t)$ is a sustainable diet consumed by sustainable practitioners in country $i$ at $t$. $P_i(t)$ is the population of country $i$ at $t$.

A diet $\mathbf{D}$ is a caloric break down of per capita consumption in 7 food groups - fruits, vegetables, grains, meats, dairy, sugar, oils. Using data from the UN FAOSTAT Food Balance Sheets, we can construct $\mathbf{D}^i(t)$, the average diet for $i$ at $t$, by adhering to the food group divisions defined in Rizvi et al. We design $\mathbf{D}^{max,i}(t)$ and $\mathbf{D}^{S,i}(t)$ by keeping caloric values under all groups in $\mathbf{D}^i(t)$ same except the caloric values under meats and dairy groups. For $\mathbf{D}^{max,i}(t)$ we replace the meats and dairy caloric values in $D^i(t)$ with the cumulative dietary consumption of countries in $I$ that consumed the most in that year in items that belong to the groups of meats and dairy. A similar approach was taken while designing $\mathbf{D}^{S,i}(t)$. Per capita meats and dairy caloric intake in $D^i(t)$ was replaced by the cumulative dietary consumption of countries in $I$ that consumed the least in that year in items that belong to the groups of meats and dairy. We take an example to explain this better. For this example we take $i$ to be the United States of America and $t$ to be 2010. From UN FAOSTAT Food Balance Sheet data, the construction of $D^{USA}(2010)$ looks as follows (all units in kcals/capita/day):

$\mathbf{D}^{USA}(2010)$ = { fruits: 122, vegetables: 163, grains: 61, meats: 621, dairy: 441, oils: 671, sugar: 591}

The meats and dairy groups, as defined by Rizvi et al. [2], contain the food balance sheet items bovine meat, goat and sheep meat, pig meat, poultry meat, eggs, milk and butter (ghee). The items bovine meat, goat and sheep meat, pig meat, poultry meat and eggs belong in the meats group and the items milk and butter(ghee) belong in the dairy group. Table 1 notes from data the highest and lowest consumers of these seven items in the year 2010 along with their respective average per capita calorie intake. We do not consider countries for which the consumption is zero while evaluating the lowest consuming countries for a concerned item. If the group values in Table 1 are added up then, the diets $\mathbf{D}^{max,USA}(2010)$ and $\mathbf{D}^{S,USA}(2010)$ would be as follows:

$\mathbf{D}^{max,USA}(2010)$ = { fruits: 122, vegetables: 163, grains: 61, meats: 1403, dairy: 749, oils: 671, sugar: 591}

$\mathbf{D}^{S,USA}(2010)$ = { fruits: 122, vegetables: 163, grains: 61, meats: 7, dairy: 6, oils: 671, sugar: 591}

Once constructed, these maximum and sustainable diets are fed into the Rizvi et al. land use evaluation model. It tells us how much land would have been spent globally if United States of America consumed these hypothetical per capita diets in 2010. We divide the land values obtained from the model with the population of USA in 2010 to receive the values of $c^{U,max}_{USA}(2010)$ and $c^{S}_{USA}(2010)$. Supplementary Figure 2 in Section 6 shows the constructed time series $c^{U,max}_i(t)$, $c^S_i(t)$ and $c_i(t)$ for USA, China, India, Russia, Brazil and Australia for every year between 1961 and 2013. Time series for Russian Federation is shown only from 1992 to 2013 since it did not exist per se prior to 1992.

#### 1.2 Data Source for $m_i(t)$ and Method for generating $m^0_i(t)$ time series

In our model, $m_i(t)$ is defined as the average income of the population of country $i$ at year $t$. We collect the per capita income data from the World Bank database [3]. We use the data listed under GDP per capita, PPP for our purpose.

In our model, $m_i^0(t)$ is defined as the minimum income required to afford the sustainable diet $\mathbf{D}^{S,i}(t)$ in $i$ at $t$. We set it to the minimum wage (and for some cases the living wage) in $i$ at $t$. We obtain the minimum wage data for the OECD countries from the Real Minimum Wage data-set compiled by the Organisation for Economic Co-operation and Development (OECD) [4]. If no minimum wage data is obtained from the first source, we set $m_i^0(t)$ to the living wage of $i$ at $t$. Living wage data is obtained from the source: [5]. If a particular country is not covered by either of the aforementioned sources, we rely on the sources compiled in the online, community maintained, article [6]. For some countries, none of the above sources provide any minimum or living wage data. These cases are specially handled and are discussed in more detail in subsection 1.4.

Since our sources of minimum wage (or living wages) data do not report these wages for all the years we wish to parameterize our model over, we evaluate $m_i^0(t)$ for all years between $t_0^i$ to $t_f^i$ using the following back-extrapolating method:

$$m_i^0(t) = m_i(t) \cdot (m_i^0(t_r^i)/m_i(t_r^i)) \quad t \in [t_0^i, t_f^i]$$

Where $t_r^i$ is the year at which minimum wage statistic was reported. Here we assume that minimum/living wage of countries maintain a constant ratio with the respective country's average income over all years in $[t_0^i, t_f^i]$. Since we have average income data for all the years in $[t_0^i, t_f^i]$, we can use the above method to back extrapolate the values of $m_i^0(t)$ for all years that lie in the period, $[t_0^i, t_f^i]$.

#### 1.3 To determine $x_{0,i}$ from $h_i$ and data

At the onset, there are four parameters unknown for the parameter estimation problem - $\kappa_i$, $\sigma_i$, $h_i$ and $x_{0,i}$. In order to reduce the dimensionality of the parameter search space, we discuss a method to evaluate $x_{0,i}$ from the first data point and $h_i$. This operation reduces the number of free parameters from 4 to 3 and assures that the $c_i$ predicted from the model at the first time point $t_0^i$ is equal to the first data point at $t_0^i$.

For a $h_i$, we can write the following at $t_0^i$:

$$L_i(t_0^i)/P_i(t_0^i) = x_i(t_0^i)c_i^S(t_0^i) + (1 - x_i(t_0^i))c_i^U(t_0^i)$$

In this equation $x_i(t_0^i)$ is the initialization, $x_{0,i}$, of the differential equation for $x_i$. $L_i(t_0^i)$ and $P_i(t_0^i)$ are taken from data. This implies that:

$$x_{0,i} = x_i(t_0^i) = \frac{c_i^U(t_0^i) - (L_i(t_0^i)/P_i(t_0^i))}{c_i^U(t_0^i) - c_i^S(t_0^i)} \tag{5}$$

Where,

$$c_i^U(t_0^i = (c_i^{U,max}(t_0^i) - c_i^S(t_0^i))(1 - e^{-h(m_i(t_0^i) - m_i^0(t_0^i))}) + c_i^S(t_0^i)$$

Hence, given a value of $h_i$, it is possible to derive $x_{0,i}$ from equation 5 provided there is knowledge about the following data at time $t_0^i$: $L_i(t_0^i)$, $P_i(t_0^i)$, $c_i^{U,max}(t_0^i)$, $c_i^S(t_0^i)$ and $m_i^0(t_0^i)$. This operation reduces the search space for the parameters from 4 to 3 as $x_{0,i}$ can be determined from $h_i$ and data implicitly.

For ease of expression in the next subsection, let us define a mapping $\xi(.)$ that relates this implicit relationship between $h_i$ and $x_{0,i}$. The function $\xi(.)$ maps the tuple $(h_i, i, t_0^i)$ to a value of $x_{0,i}$ such that equation 5 is respected. That is,

$$x_{0,i} = \xi(h_i, i, t)$$

It is worth noting that equation 5 also puts bounds on the search space of $h_i$ since $x_{0,i}$, by definition, always lies between 0 and 1. Bounds on $h_i$ makes further reduction in the parameter search space.

#### 1.4 Algorithm for Parameter Estimation

Note from equation 1 that $c_i(t)$ is also function of the parameters $\kappa_i$, $\sigma_i$, $h_i$ and $x_{0,i}$. We use the following alternate representations of $c_i(t)$:

$$c_i(t) \equiv c_i(t, \kappa_i, \sigma_i, h_i, x_{0,i}) \equiv c_i(t, \kappa_i, \sigma_i, h_i, \xi(h_i, i, t)) \equiv c_i(t, \kappa_i, \sigma_i, h_i)$$

Given the time series of per-capita land use of a country $i$ in $[t_0^i, t_f^i]$ as input data, the task is to find the optimal parameters $\kappa_i, \sigma_i$ and $h_i$ so that the loss function defined between the data and the model is minimized.

The loss function that we choose to determine the deviation of the model output from the data is a weighted root mean square metric where the weights are linearly increasing from $t_0^i$ to $t_f^i$. Imagine $N^i$ time series data points

$\{d_1, d_2, ..., d_{N^i}\}$ and a corresponding model output time series $\{m_1(\theta), m_2(\theta), .., m_{N^i}(\theta)\}$. We define our weighted loss function $L(\theta)$ between data and model output as follows:

$$L(\theta) = \frac{1}{N^i \sum_{k=1}^{k=N^i} w_k} \sum_{k=1}^{k=N^i} w_k (d_k - m_k(\theta))^2, \quad w_k = k \tag{6}$$

Let us represent the time series data that we have of per capita land use for $i$ from $t_0^i$ to $t_f^i$ as: $\{c_i^d(t_0^i), .., c_i^d(t_f^i)\}$. For each country, $i \in I$, we have $N^i$ equi-spaced data points including $t_0^i$ and $t_f^i$ . That is $N^i = t_f^i - t_0^i + 1$. Since we deal with yearly data, $t_0^i$ and $t_f^i$ are the starting and ending years. Our parameter estimation algorithm tries to evaluate the optimal values of $\kappa_i$, $\sigma_i$ and $h_i$ so that the following is minimized:

$$\frac{1}{N^i \sum_{t_j \in [t_0^i, t_f^i]} w_{t_j}} \sum_{t_j \in [t_0^i, t_f^i]} w_{t_j} (c_i^d(t_j) - c_i(t_j, \kappa_i, \sigma_i, h_i))^2, \quad w_{t_j} = t_j - t_0^i \tag{7}$$

For parameter estimation, we do not use the closed loop feedback. That is, we do not feed equation 3 into equation 4. Instead, we use the time series of total global land use, $L^G(t)$, into equation 4 (evaluated from data). This time series is the same as the yellow time series seen in Figure 1 of this text. This time series is evaluated using the model described in [2]. This model is available as Python scripts in: Saptarshi07/Dietary-Trends-Tools.

For countries where the minimum/living wage statistic is absent completely, we optimize for parameters $\sigma_i$, $h_i$ and $m_i^0(2018)$ while assuming a value of $\kappa_i$. The assumed value of $k_i$ is estimated by averaging the $\kappa$ values of geographically nearby countries. For example, two such countries are Norway and Sweden. We use the evaluated value of $\kappa$ from Denmark and Finland to fill in the value of $\kappa$ for Norway and Sweden. Similarly, for Niger, we take the evaluated $\kappa$ values of Mali, Nigeria and Benin to fill in the $\kappa$ value for Niger.

We use a combinatorial optimization approach in our numerical algorithm to determine the near-optimal parameter values for our model. The three parameters that we estimate can either be $\kappa$, $\sigma$ and $h$ or, they can be $\sigma$, $h$ and $m^0(t_r^i = 2018)$ depending on whether minimum wage statistic is available for the country. Let us denote the three parameters we want to estimate with our algorithm as $\theta_1, \theta_2$ and $\theta_3$.

At the beginning of our algorithm, we set theoretical bounds for our parameters. The range for parameter $\theta_i$ is denoted as $[\theta_{i,l}, \theta_{i,u}]$. Here $l$ and $u$ stand for lower and upper bounds. For example, the theoretical bounds for $log(\kappa)$ was taken to be $[-12, 0]$ and the theoretical bounds for $log(\sigma)$ was taken to be $[0, 12]$. For $log(h)$ it was $[-8, 0]$ and for $m^0(t_r^i)$ it was $[0, m(t_r^i)]$. All logarithms here are with base 10. The search for parameters is conducted in the logarithmic space. The bounds for parameters remain constant for every country throughout the parameter estimation process. Our numerical algorithm for optimization requires two other input hyper-parameters. We call them the depth $(D)$ and runs $(R)$ of the algorithm.

**Algorithm:**

1. Get R equidistant points inside each bound range, $[\theta_{i,l}, \theta_{i,u}]$. Let these points be - $\{\theta_{i,0}, .., \theta_{i,k}, ...\theta_{i,R-1}\}$. Here $i = 1, 2, 3$. $\theta_{i,l} = \theta_{i,0}$ and $\theta_{i,u} = \theta_{i,R-1}$

2. For all $R^3$ combination of parameters, find the combination for which $L(\theta)$ is minimum. Let them be $\theta_{1,a}$, $\theta_{2,b}$ and $\theta_{3,c}$.

3. Reset the bound for parameter 1 as follows: $\theta_{1,l} = \theta_{1,a-1}$ and $\theta_{1,u} = \theta_{1,a+1}$. If $a = 0$ then $\theta_{1,l} = \theta_{1,a}$ and if $a = R - 1$ then $\theta_{u,l} = \theta_{1,a}$. Reset bounds for parameters $\theta_2$ and $\theta_3$ similarly.

4. Do step 1-3 for remaining $D - 1$ (depth - 1) number of times.

5. The optimal evaluated parameters are $\theta_{1,a}$, $\theta_{2,b}$ and $\theta_{3,c}$ after $D$ runs.

We use $D = 5$ and $R = 10$ for the parameter estimation process. We use the loss function defined in Equation 7 as $L(\theta)$. As mentioned earlier, the search is conducted in the log-space. This means that we obtain the optimal parameters in their logarithmic form - $log(\theta_{1,a}), log(\theta_{2,b})$ and $log(\theta_{3,c})$.

#### 1.5 Choice of $t_0^i, t_f^i$ and discussion

As mentioned in the previous subsection 1.4 all the time series data that we work with are yearly and between 1961 to 2013. So, $t_0^i$ and $t_f^i$ are also years between 1961 and 2013. For every country $i$ a decision needs to be made before parameterization about the choice of $t_0^i$ and $t_f^i$. The following points note down the steps for this choice:

1. $t_0^i$ and $t_f^i$ are always integer values in $[1961, 2013]$. $t_0^i < t_f^i$ should always be maintained.

2. For no year in $[1961, t_0^i) \cap (t_f^i, 2013]$ per capita land use data for $i$ should be available.

An example of $i$ where $t_0^i$ is not 1961 is the Russian Federation. For Russia, $t_0^i$ is 1992 since the country did not exist by that name prior to 1992 and so no data for per capita land use is available for it between 1961 and 1991. Similarly, an example where $t_f^i$ is not 2013 is the USSR.

### 2 Brief description of the land use evaluation model in [2]

We represent the Rizvi et al. model as a function, $R(.)$, that maps a diet $\mathbf{D}$, a country $i$, and a year $t$ into a land use value. That is, if the population of $i$ in year $t$ consumed the average per capita diet $\mathbf{D}$, $R(\mathbf{D}, i, t)$ hectares of land would have been spent, globally, to generate the demand. For this function $t$ is an integer such that $1961 \leq t \leq 2013$. A diet is defined, mathematically, as a column vector of length 7. Numeric value of the vector components represent daily caloric intake in the food groups of fruits, vegetables, grains, meats, dairy, oils and sugar. For every item in the food balance sheet (that is assigned a parent food group), data for food supply quantity (in kilograms per capita per day) and food supply (kcals per capita per day) is provided simultaneously for a country at a year. This helps in evaluating the energy to mass conversion factor for a food item $j$ in a country $i$ at a year $t$. Let $k$ be a food group and $I_k$ be the set of items listed under the food group $k$. We define the set of food groups as $G$ (and $k \in G$). $D_k$ is the per-capita daily calorie intake of food group $k$, as defined by the diet $\mathbf{D}$. We represent the food supply data for an item $j$ in $i$ at $t$ as $f_j^{i,t}$. Then, the per-capita calorie intake of an item $j \in I_k$, in $i$ at $t$, $d_j^{i,t}$, can be evaluated as the following:

$$d_j^{i,t} = D_k \cdot (f_j^{i,t} / \sum_{j \in I_k} f_j^{i,t})$$

If the food supply quantity of an item $j$ in $i$ at $t$ be represented as $s_j^{i,t}$, then the kilo calorie to kilogram conversion factor in $i$ at $t$, $c_j^{i,t}$, can be evaluated as follows:

$$c_j^{i,t} = \frac{s_j^{i,t}}{f_j^{i,t}}$$

The yearly mass demand, $R_j^{i,t}$ of item $j$ in $i$ at $t$, in tonnes, would then be:

$$R_j^{i,t} = d_j^{i,t} \cdot P^{i,t} \cdot 365 \cdot \frac{c_j^{i,t}}{1000}$$

Where $P^{i,t}$ is the population of the country $i$ in year $t$. Note that here we have made the following assumption: for any arbitrary dietary intake of a food group $k$ (say $D_k$), the distribution of $D_k$ across the group items maintains the same proportion to that of the reported data. That is, if the average caloric intake of bovine meat in USA in 1980 was $1/4^{th}$ of the total calorie intake of meats (let's say 500 kcals/capita/day), then any other dietary intake, $D_k$, for meats, would have $1/4^{th}$ of it dedicated to bovine meat (in USA, in 1980).

Now, we define another conversion factor $C_j$ called the source conversion factor for a food item $j$. The source conversion factor is independent of the country or the year (hence it does not have the superscripts $i$ and $t$). A source conversion factor converts the mass of a food item to an equivalent mass of its source item. For most items this conversion factor is 1. However, for items like beer, wine, butter etc, the value is not unity. For example, source of beer is barley and its mass conversion factor is 4.78. To generate 1 tonne of beer, 4.78 tonnes of barley is required, on average.

The food balance sheet reports data for the Domestic Supply Quantity (in tonnes) and the Import Quantity (in tonnes) of every food item $j$ into a country $i$ at a year $t$. The Import Quantity data element, $I_j^{i,t}$, indicates the amount of the food item $j$ that was imported into $i$ in year $t$. The Domestic Supply Quantity data element, $D_j^{i,t}$, indicates the amount of $j$ that is available to the population of $i$ at $t$ for domestic utilization. The ratio of Import Quantity to Domestic Supply Quantity is defined as the import dependency ratio, IDR, of $j$ in $i$ at $t$ - $IDR_j^{i,t}$. That is,

$$IDR_j^{i,t} = I_j^{i,t} / D_j^{i,t}$$

The quantity $j$'s source that comes in through import to meet the dietary demand of $i$ at $t$ is then,

| Country Name ($i$) | Start Year $t_0^i$ | End Year $t_f^i$ | $m_i^0(2018)$ (USD) | $log_{10}(\kappa_i)$ | $log_{10}(\sigma_i)$ | $log_{10}(h_i)$ | $x_{0,i}$ |
|---|---|---|---|---|---|---|---|
| United States of America | 1961 | 2013 | 18262 | -11.98 | 10.09 | -2.61 | 0.65 |
| China | 1961 | 2013 | 3600 | -11.88 | 10.20 | -2.30 | 0.74 |
| India | 1961 | 2013 | 1500 | -12.00 | 9.82 | -0.1 | 0.79 |
| Russian Federation | 1992 | 2013 | 4734 | -11.34 | 8.00 | -3.03 | 0.68 |
| Brazil | 1961 | 2013 | 5114 | -11.77 | 10.20 | -1.95 | 0.86 |
| Australia | 1961 | 2013 | 12600 | -11.52 | 0.007 | -1.61 | 0.58 |

Table 2: Model fitting results for six countries of the world. Per capita land use model is fitted to data here.

$$I_{j,F}^{i,t} = \frac{IDR_j^{i,t} R_j^{i,t}}{C_j}$$

Similarly, the quantity $j$'s source that comes from within the borders of $i$ to meet the dietary demand of $i$ at $t$ is given by:

$$D_{j,F}^{i,t} = \frac{(1 - IDR_j^{i,t}) \cdot R_j^{i,t}}{C_j}$$

The units of $D_{j,F}^{i,t}$ and $I_{j,F}^{i,t}$ are in tonnes (per year). $Y_j^{i,t}$ and $\bar{Y_j}^t$ are defined as the yield of source of $j$ in $i$ and the average yield of source of $j$ in the world respectively in $t$. Methods for calculating these are not included in this Supplementary Information document. For more information on it we advise the reader to check the Supplementary Information of [2]. The units of these variables are in tonnes per hectare. Then, the expected land required globally to produce the demand for item $j$ in $i$ at $t$ is:

$$L_j^{i,t} = \frac{D_{j,F}^{i,t}}{Y_j^{i,t}} + \frac{I_{j,F}^{i,t}}{\bar{Y_j}^t}$$

Hence the total global land required to produce the average dietary demand $\mathbf{D}$ for a country $i$ in $t$ is:

$$L^{i,t} = \sum_{k \in G} \sum_{j \in I_k} L_j^{i,t}$$

### 3 Results of parameter fitting

The results for the parameter fitting process for six select countries are summarized in table 2. Figures 3 and 4 respectively show the model fits in the per-capita scale and the total land-use scale.

In Figures 5, 6, 7 we see a global heat map for the rescaled parameters $\kappa_i$, $\sigma_i$ and the parameter $h_i$ for 166 of the currently existing countries in the world. The countries in colour grey are the ones for which parameters are not estimated. Figure 8 is the global heat map for the ratio $m_i^0(t_r^i)/m_i(t_r^i)$. $t_r^i$ is the year when living wage or minimum wage was reported. Note that these heat-maps show estimated parameters in their logarithmic form, i.e in log base 10 scale. In section 4 we list the countries that were chosen for our analysis. The reason for inclusion and exclusion of countries in the analysis are also explained in section 4.

At 2013, we achieve 1.24 % error, globally, from our model. That is, summed over the 166 countries we estimate parameters for, our model output is 1.24 % deviated from the global land use data in 2013.

In Figure 12, absolute model errors are shown year-wise in a box-whisker plot. Statistics of absolute percentage error of the country-level model output relative to data is plotted. At 1961, error is zero due to our parameterization procedure (see Section 1.4). Average absolute errors are always bounded between 0 % to 10 %. At 2013, average absolute percentage error is around 5% but global error is 1.24% as errors of several countries cancel each other.

### 4 Countries included in the analysis and discussions

A total of 180 countries are listed by the FAO Food Balance Sheet, 2013 [1]. These consist of both currently existing countries and countries that have ceased to exist (e.g USSR). Out of these 180 countries, parameter fitting was done for 166 countries - all of which are currently existing. The countries that were excluded because they no longer exist are - USSR, Yugoslavia SFR, Former Sudan, Ethiopia PDR, Serbia and Montenegro, Czechoslovakia, Netherlands Antilles and Belgium-Luxembourg. Currently existing daughter nations of these formerly existing countries are fitted for parameters instead. The DPRK is excluded because our source does not provide any food consumption data for it. Since our sources provide no per capita income data for Taiwan, it was eliminated for parameter estimation process too. The countries Saint Kitts and Nevis, Bermuda, Dominica, Antigua and Barbuda, Kiribati, Grenada and Saint Lucia are excluded because they have no model projections for population and income till 2100. OECD Env-Growth [7] is one of the models that is complied in the IPCC SSP Projection database for population and income. Since we use the projections for per capita income and population from this model in our land use projection model, the aforementioned seven countries are excluded.

The time series for total global land use due to consumption (evaluated as the yellow time series in Fig 1) was calculated yearly for all years between 1961 to 2013. It accounts for land consumed by all 180 countries minus Taiwan and Yugloslavia SFR. This calculation is done using the model in [2] and it accounts for the periods of existence of nations.

In Figures 5, 6, 7 and 8 we see global heat maps for the parameters of 166 nations. The countries for which parameters are not estimated are coloured in grey. Note that countries such as Papua New Guniea, DRC, Somalia, Syria and Libya are marked grey because consumption data for them are not reported by the UN FAOSTAT Food Balance Sheet, 2013 [1]. That is, they do not belong to the initial set of 180 countries.

We project for 164 countries out of the 166 countries that have been parameterized. The countries that are eliminated from the projection analysis are French Polynesia and New Caledonia.

### 5 Methods for projection analysis

Land use projections are made till the year 2100. All projections begin from 2011. We provide country level land use projections yearly between the start and end year of projections. We project for 164 countries. We call this set of 164 countries $I$. Details about their choices are explained in Section 4. We choose 2011 as the start year because all the 164 countries from our set exist as nations thereafter. One of the primary requisites for land use projection is the availability of country level scaled down projections of population and income. We use five population and income scenarios in our projection analysis. These scenarios have been pre-defined in [8] and are popularly known as the IPCC SSP scenarios. All of the five scenarios under SSP (SSP1-5) have country level population and income projection through an ensemble of models. For our purpose, we use the projections under the model OECD Env-Growth [7]. The projections for population and income are available from 2010 to 2100 at an interval of 5 years. We use spline interpolation of order 3 to interpolate projected values of income and population between the 5 year intervals. Story line description of these SSP scenarios are available in [8].

Another requisite for projection analysis is the projection of the time series $c^{U,max}$ and $c^S$ for every country $i$ in our set $I$ till 2100. We construct scenarios for the qualitative evolution of these time series. These scenarios are defined by a continuous variable $f$ called *future yield variable*. It lies between 0 and 1 and is represented by $f$. In the following subsection we discuss the method for constructing the consumption scenarios.

#### 5.1 Future yield, $f$, scenarios

Let $c$ be the concerned time series, defined between 1961 to 2013, that we wish to project till 2100. The series $c$ can either be $c^{U,max}$ or $c^S$ for a country $i$. First, we fit an exponential of form $y = ae^{bt}$ to a truncated $c$ series. This truncated version of $c$ is the time series of $c$ from 1990 to 2013. If $b < 0$ we call the series trend is decreasing and if $b > 0$ we call the series trend increasing. Here, $a$ and $b$ are constants. We extrapolate the time series $c$ till 2100 (starting from 2013 onward) using the following equations:

$$c(t) = \begin{cases} c(2013) - (c(2013) - c(2013)f)(1 - e^{-\beta(t-2013)}), & \text{if initial trend is decreasing} \\ c(2013) + c(2013)f(1 - e^{-\beta(t-2013)}), & \text{if initial trend is increasing} \end{cases}$$

Here $f$ is the tune-able parameter - a real number between 0 and 1 that defines the future yield scenario. Note that for the above equation $t > 2013$. The exponent $\beta$ is adjusted such that continuity is maintained at 2013 between the initial trend, $ae^{bx}$, and the projected trend $c(t)$. That is,

$$\beta = \begin{cases} -\frac{1}{c(2013)} \frac{abe^{2013b}}{1-f}, & b < 0 \\ \\ \frac{abe^{2013b}}{c(2013)f}, & b > 0 \end{cases}$$

In Figure 9, we show two examples of $c^{U,max}$ and $c^{S}$ projection till 2100 using the above method. The two countries that are chosen are USA and Netherlands. USA shows a decreasing initial trend whereas Netherlands shows a initial increasing trend.
Intuitively, if the trend is decreasing, a $f$ scenario implies that both $c^{U,max}$ and $c^{S}$ can, at lowest, be $f$ times their 2013 value. Similarly, for an increasing trend, a $f$ scenario means that $c^{U,max}$ and $c^{S}$ can be at most $1+f$ times their 2013 value. The rates at which they approach these bounds are determined by their historical trend from 1990 to 2013.

#### 5.2 Projection Methods, Continents and Parameter Planes

Mathematically, the projection model is the coupling of the country level model of all the countries $i$ in $I$. That is, equation 1, 2, 3 and 4 taken together for all $i$ in $I$ represent our projection model. Equations 1, 2 and 4 of every $i$ is coupled to all other countries in $I$ through equation 3. We start projecting from 2011 and continue till 2100 while making yearly projections.

We work with 5 continents in our work. They are Africa, Americas, Asia, Europe and Oceania. North, Central and South America is clubbed together into one as the Americas. There are 45, 30, 43, 38 and 6 countries in these 5 continent groupings respectively. Russia is considered to be in Europe. Turkey is considered to be in Asia.

In the parameter planes that we analyze in the main text, we vary $\kappa$, $\sigma$ and $h$ from their baseline values. For all of the parameter planes, we make increments to parameters by -100% to 200% of their baseline values.

#### 6 Supplementary Figures

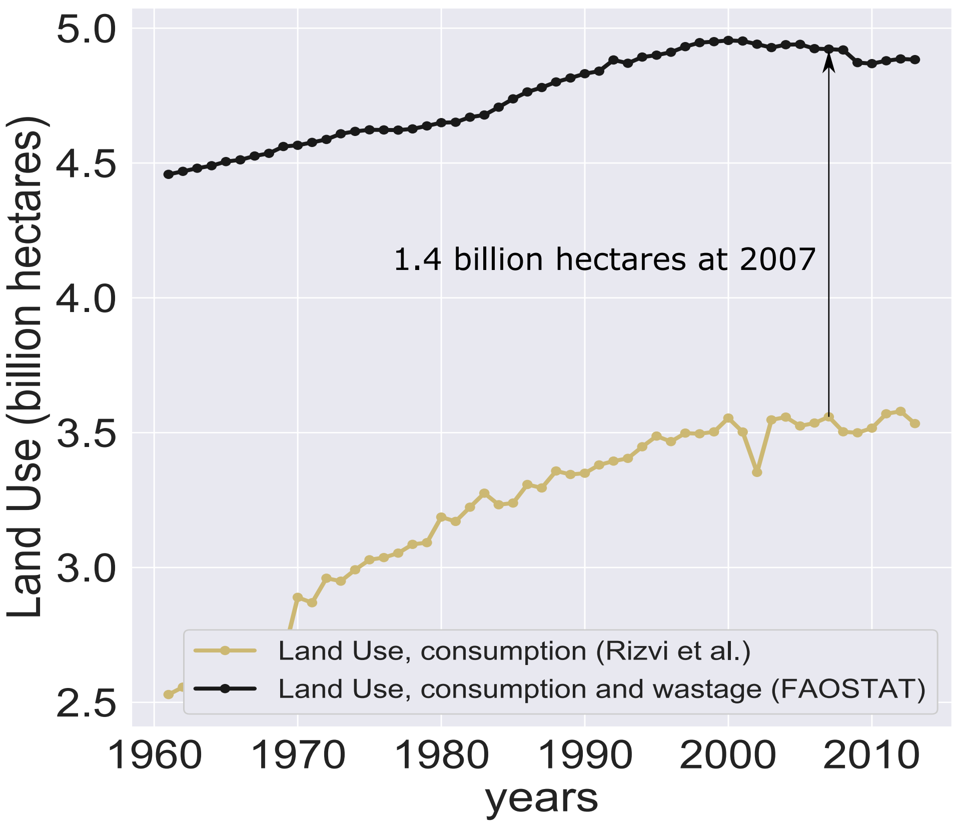


Figure 1: Time Series data for Global Agricultural Land Use from 1961 to 2013. The yellow series is the data generated by the model in [2]. It accounts for the land that was spent for generating the food for human consumption. The black series is the data collected from [1]. It accounts for the land that was spent to generate the consumed food and the wasted food. We use the annual time series in yellow as $L^G(t)$ in the parameter estimation process.

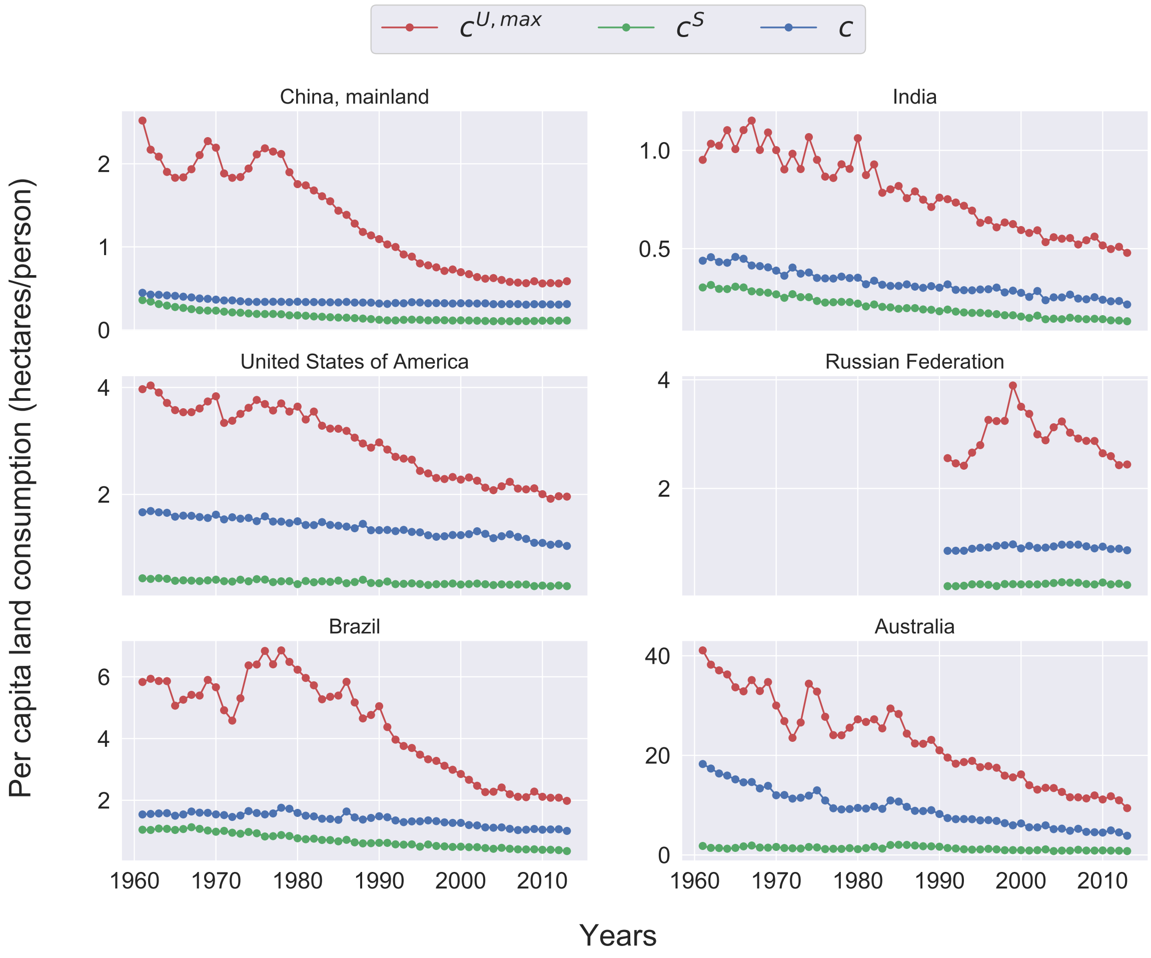


Figure 2: The constructed time-series of $c^{U,max}(t), c^S(t)$ and $c$ for United States, China, India, Russian Federation, Brazil and Australia using the method described in Supplementary Section 1.1. $c^{U,max}$ is the maximum upper limit of per capita land use in a country. $c^S$ is the per capita land use of sustainable practitioners in a population. $c$ is the average per capita land use of a population. Time series are calculated using method in Rizvi et al. [2]. See Section 1.1 for details. Note that results for Russian Federation are only shown for years between 1992 and 2013 because it did not exist by that name prior to 1992.

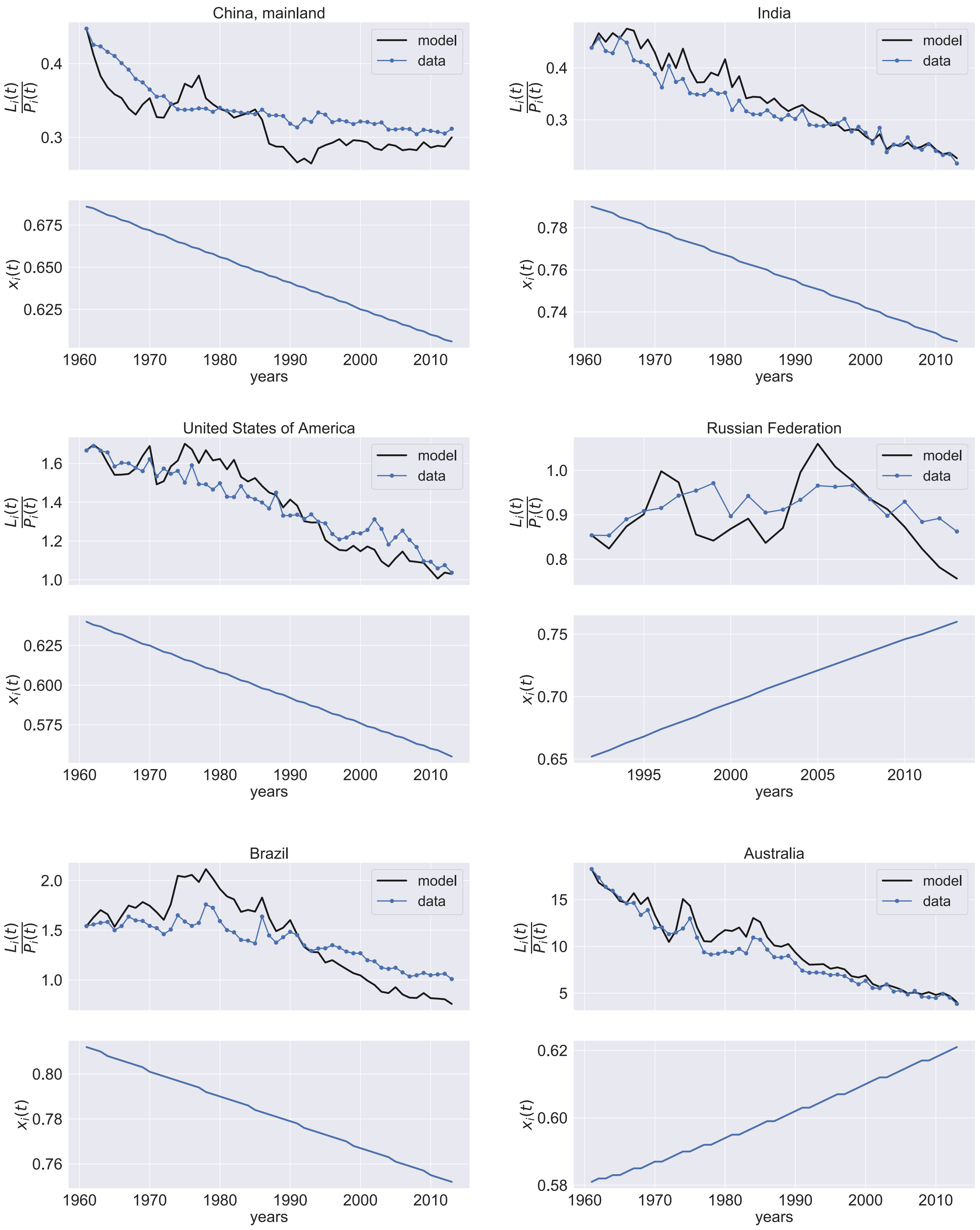


Figure 3: Model fitted estimation and data for per capita land use in six countries along with model fitted estimation of fraction of sustainable diet practitioners over the time span between 1961 and 2013. Results for Russian Federation are shown from 1992 onwards since it did not exist prior to that year. The units of $L_i(t)/P_i(t)$ are in hectares per person.

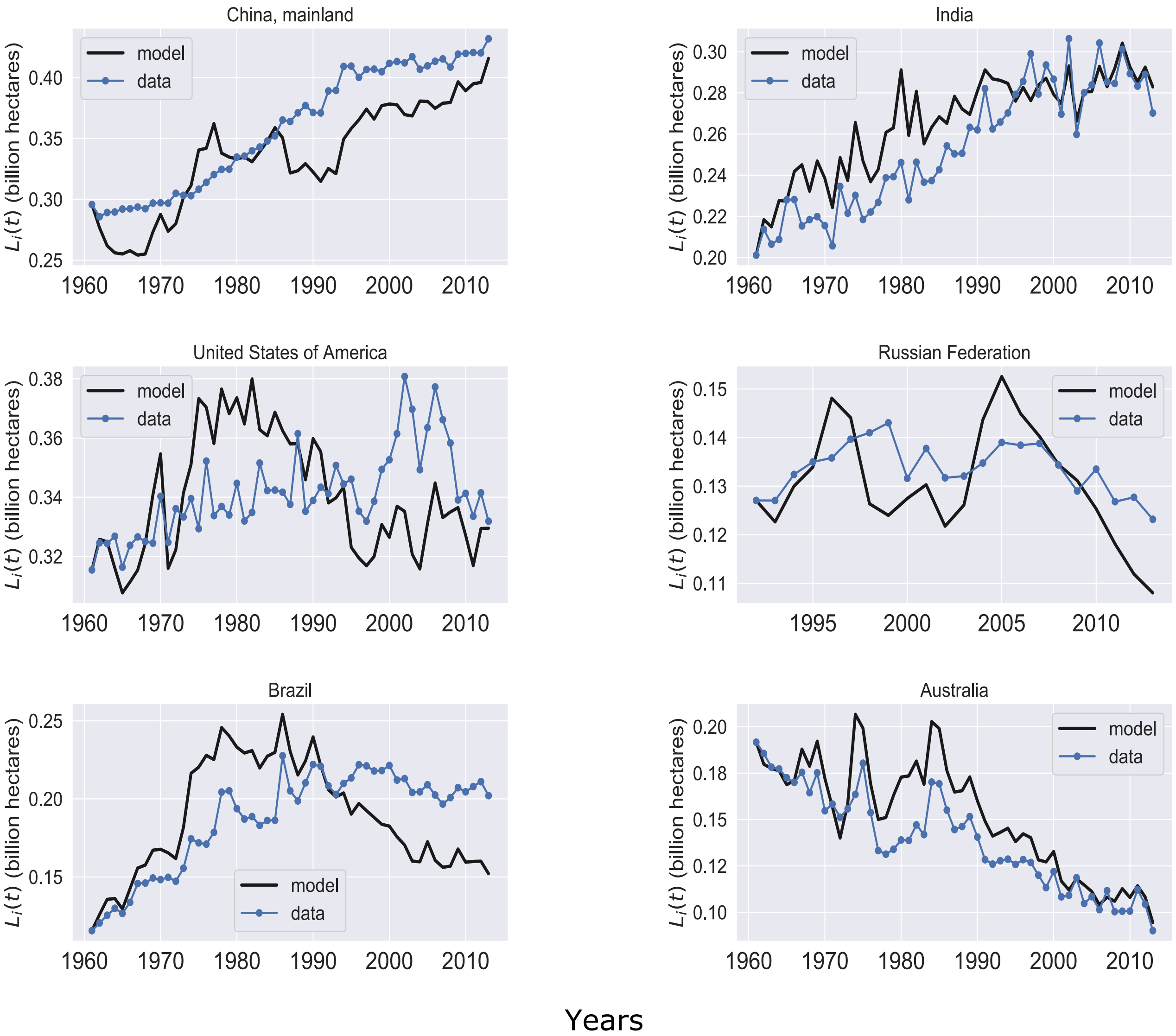


Figure 4: Model Prediction and Data for total land consumed $L_i(t)$ due to food demand of countries for years from $t_0^i$ to 2013. For all countries here, except Russian Federation, $t_0^i$ is 1961. For Russia, $t_0^i = 1992$. The y-axes units in these plots are billions hectares.

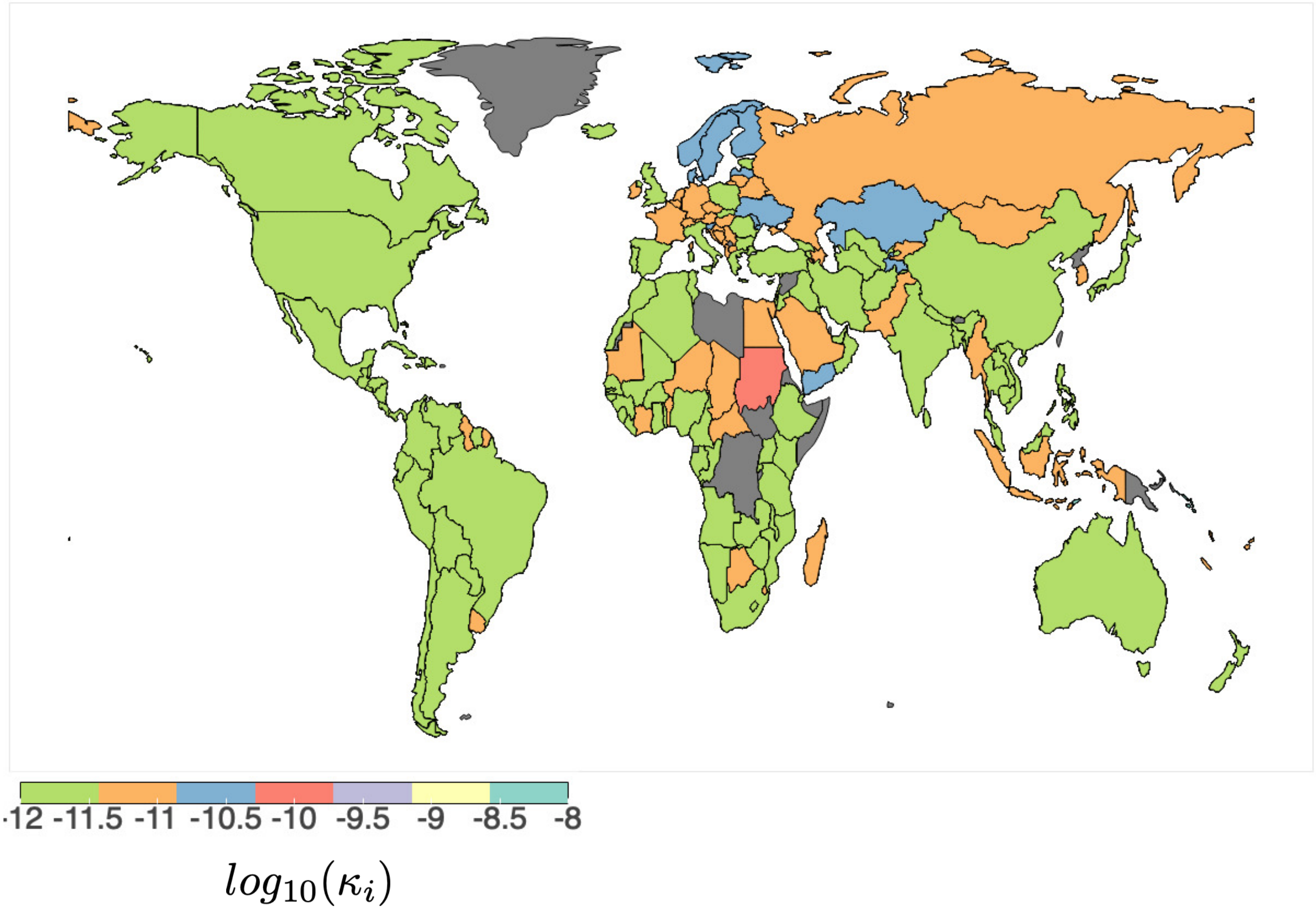


Figure 5: Global heat map of $log_{10}(\kappa_i)$ values for 166 countries in the world. The countries in grey have not been estimated for parameters.

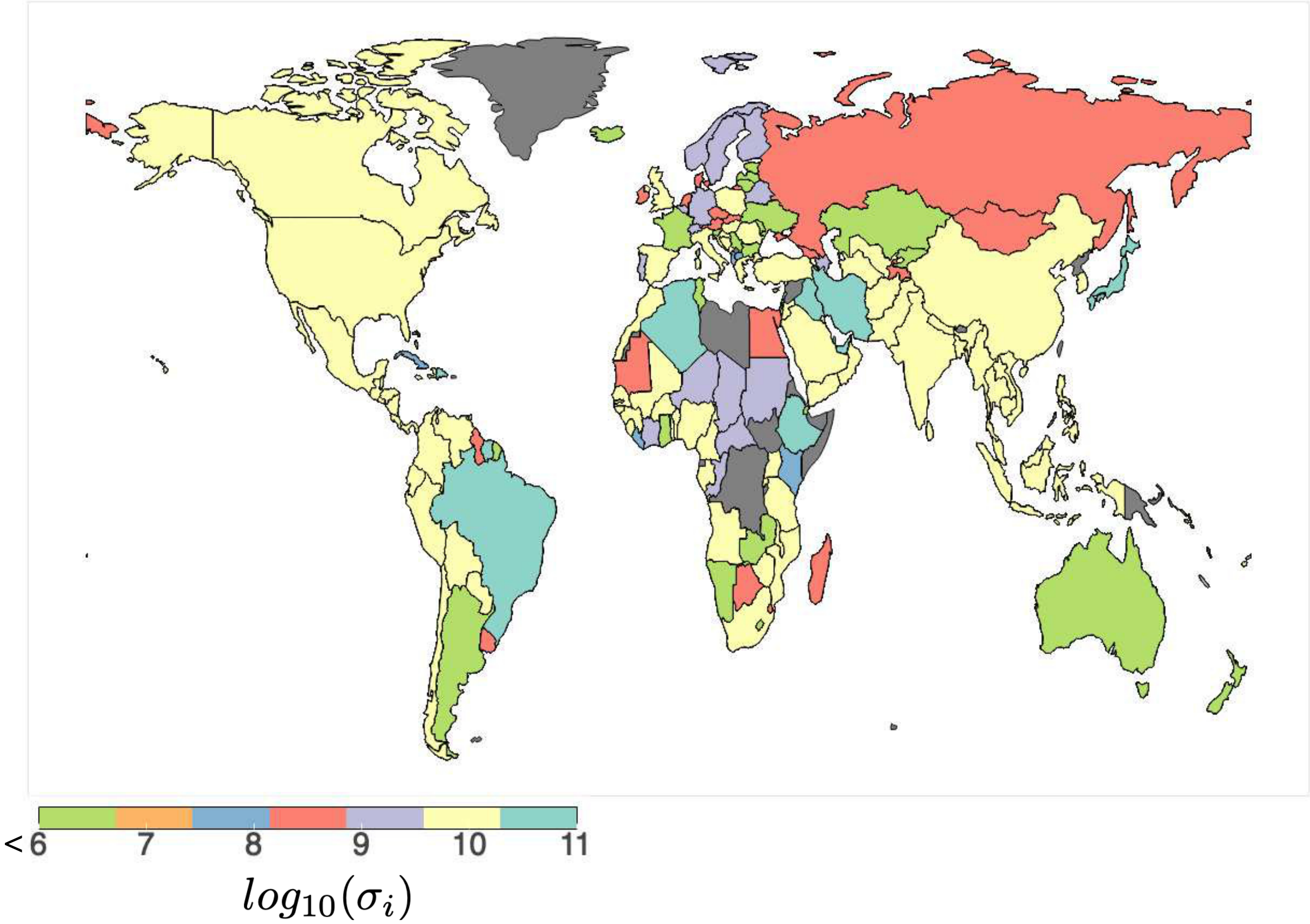


Figure 6: Global heat map of $log_{10}(\sigma_i)$ for 166 countries in the world. The countries in grey have not been estimated for parameters.

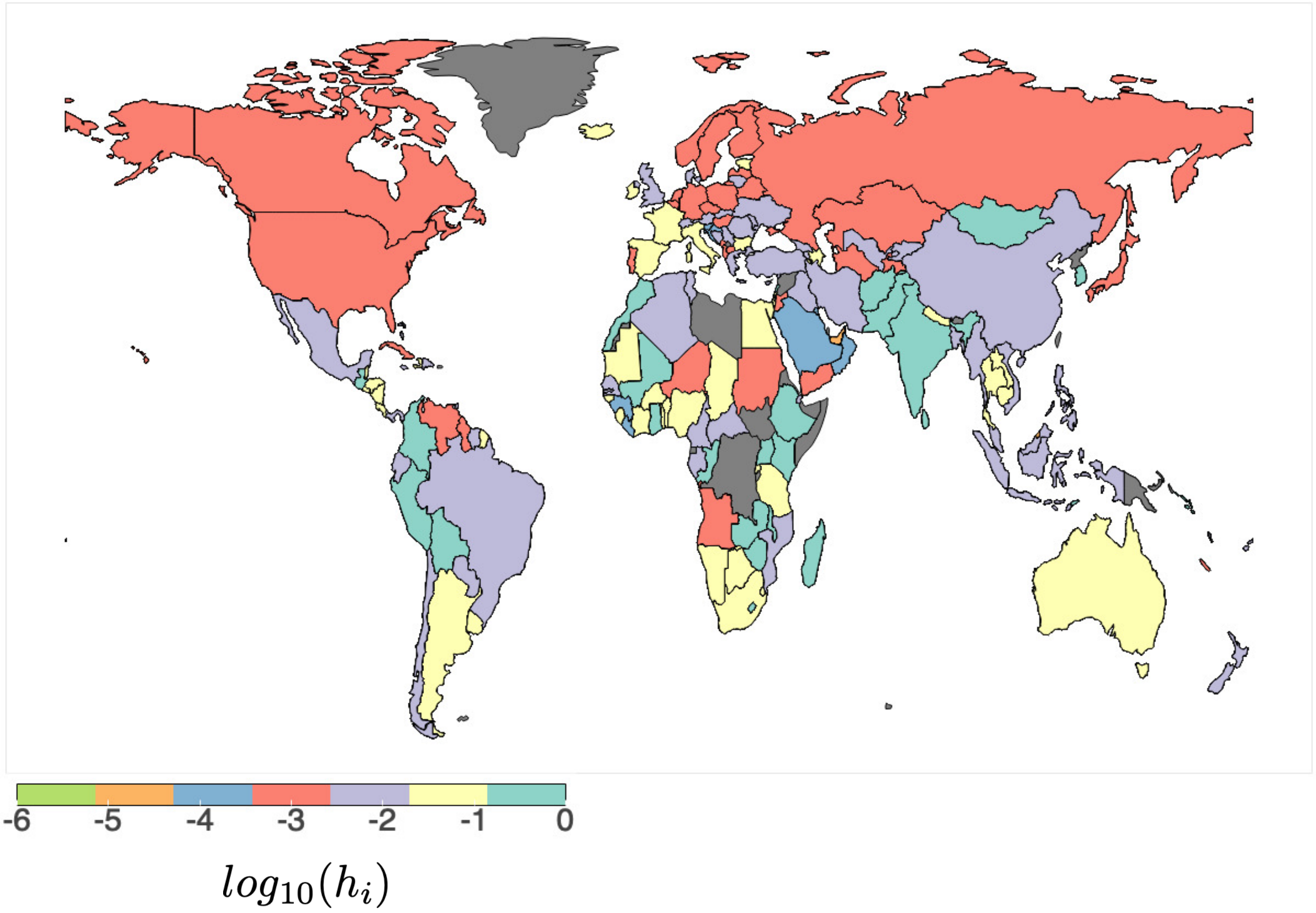


Figure 7: Global heat map of $log_{10}(h_i)$ for 166 countries in the world. The countries in gray have not been estimated for parameters.

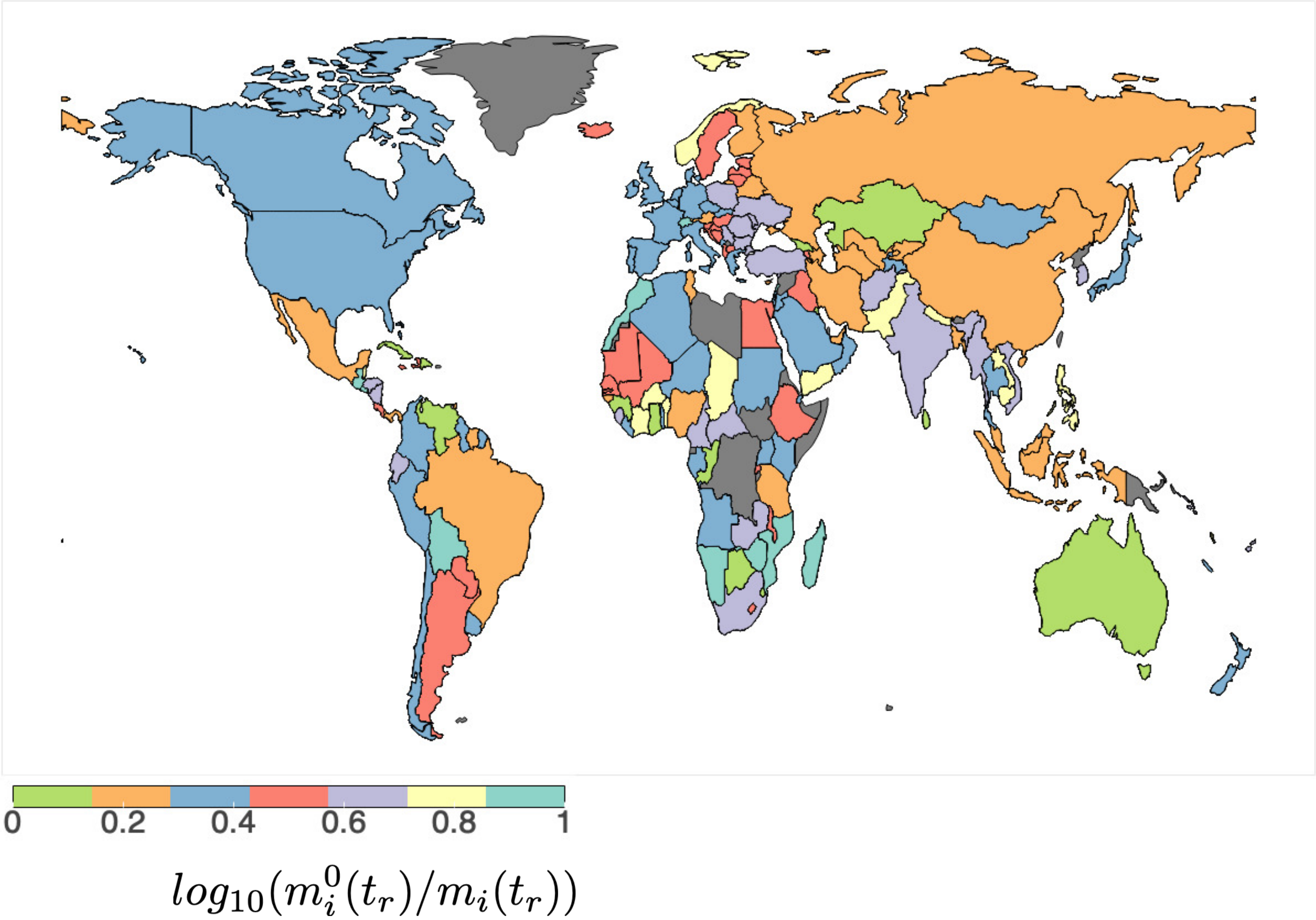


Figure 8: Global heat map of $m_i^0(t_r^i)/m_i(t_r^i)$ for 166 countries in the world. $t_r^i$ is the year when living-wage/minimum wage $m^0$ was reported. $m_i(t_r^i)$ is the average income of country $i$ in $t_r^i$.

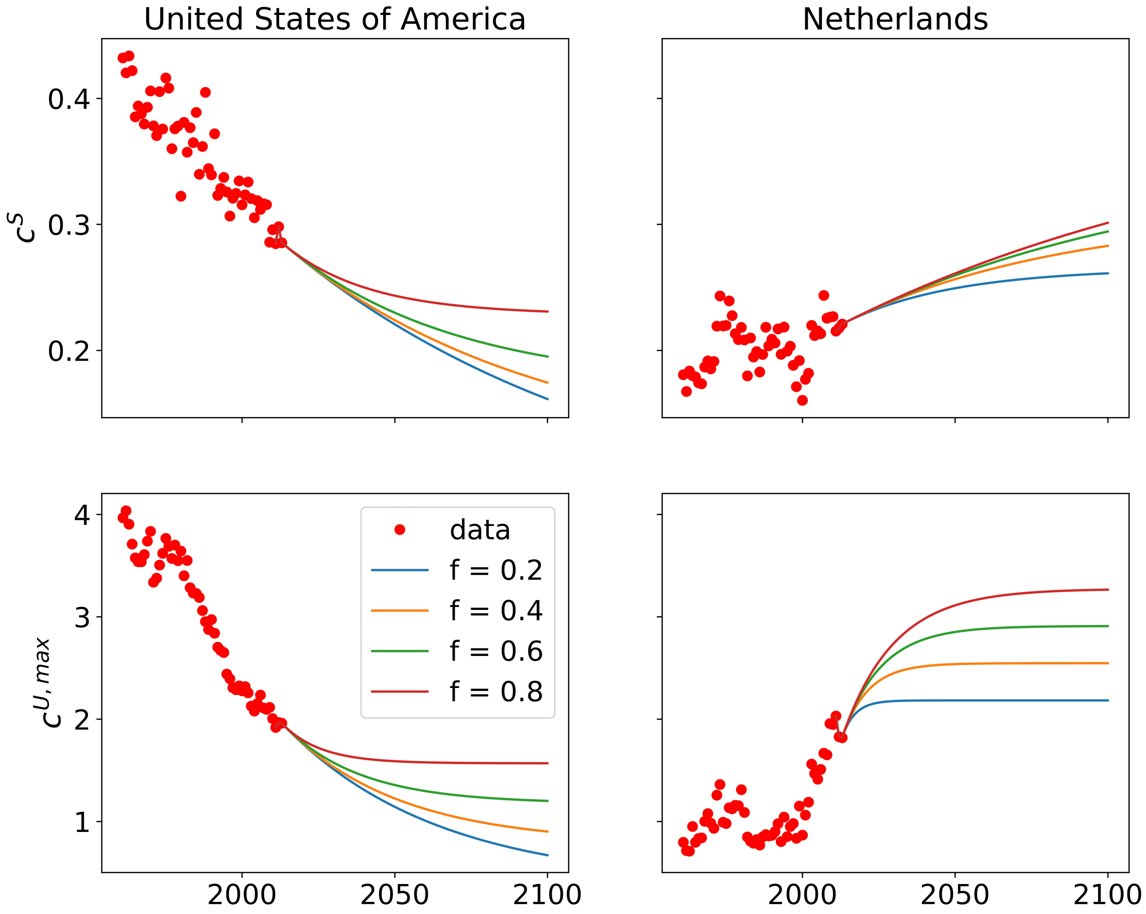


Figure 9: Projections of $c^{U,max}$ and $c^S$ for USA and Netherlands till 2100 under future yield, $f$ scenarios. USA has an initial decreasing trend while Netherlands has an initial increasing trend. In this plot, results for $f$ scenarios 0.2, 0.4, 0.6 and 0.8 are shown.

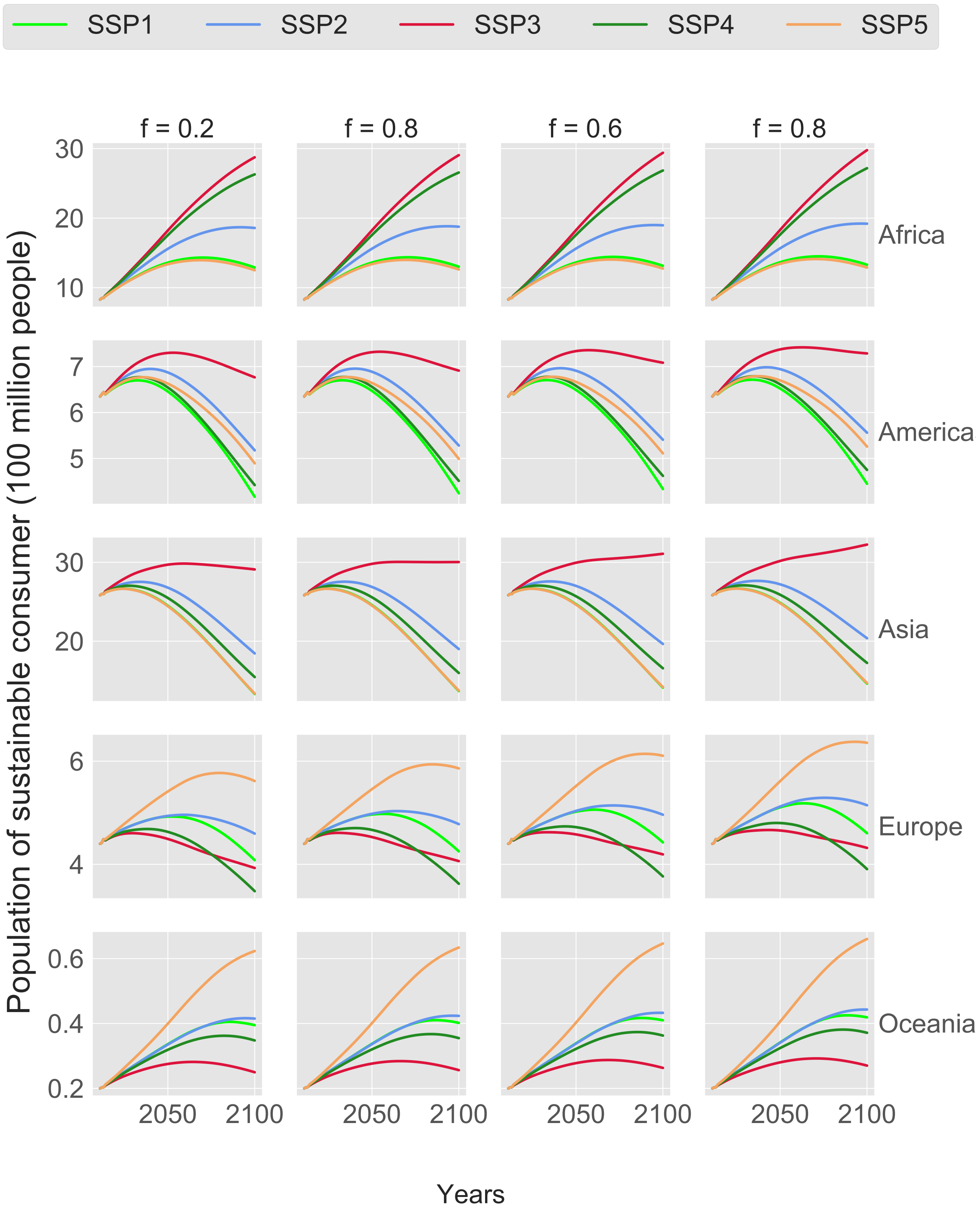


Figure 10: Model projections for number of people consuming sustainably in the 5 regions from 2011 to 2100 under 20 scenario combinations. At the continental level, increase (or decrease) in sustainable consumer fraction does not imply increase (or decrease) in the population of people consuming sustainably.

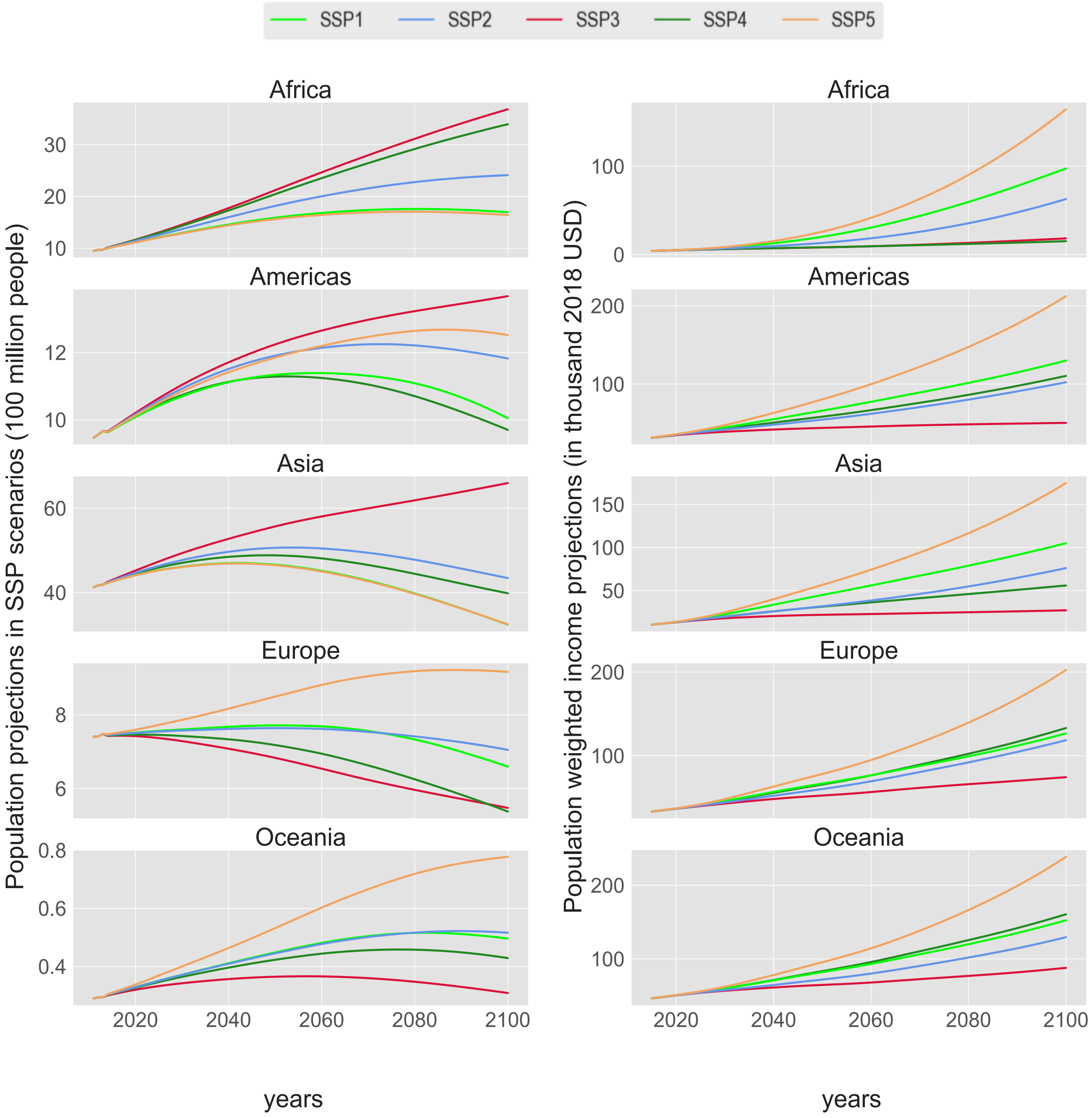


Figure 11: Population and income projections under SSP scenarios for five distinct geographical regions. Income projections are calculated by taking population weighted average of country level projections. SSP3 shows lowest income growth while SSP5 shows the highest. For Europe and Oceania, SSP3 shows lowest population growth whereas it shows highest population growth for Africa, Asia and the Americas. Segregation of continents (regions) are done based on the FAOSTAT database.

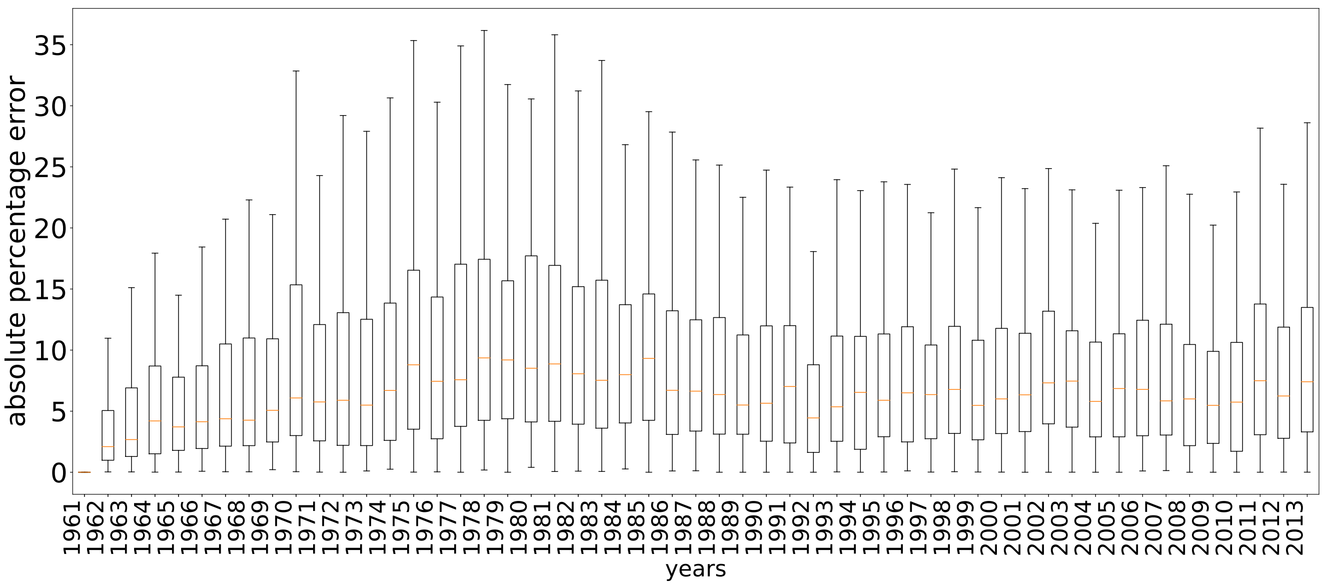


Figure 12: Box and whisker plot showing absolute error of country-level model output with respect to data over the years from 1961 to 2013. Average absolute errors always remain below 10%.

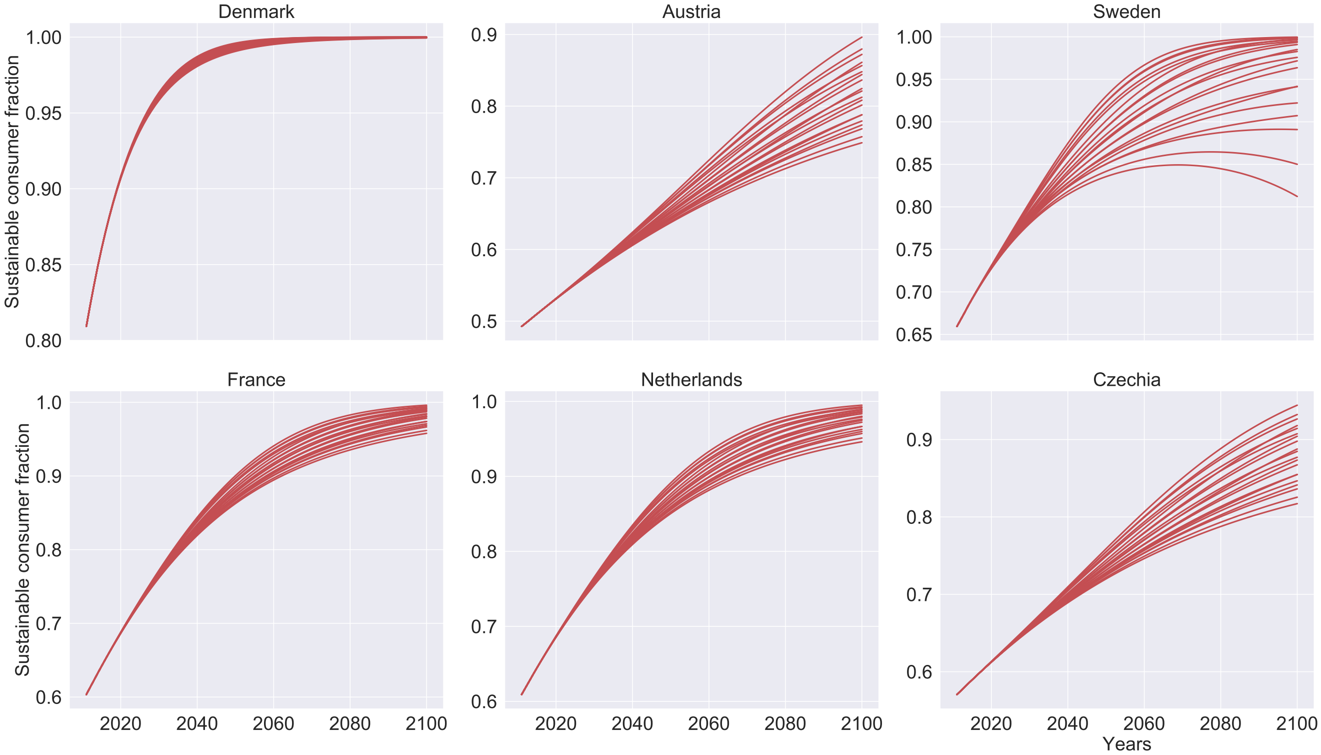


Figure 13: Model projection of sustainable consumer fraction in six European countries that show relative increase in sustainable proportion in their population between 2013 and 2100. Results are shown for 25 scenario combinations (unlabeled).

[4] Organisation for Economic Co-operation and Development. Real minimum wages, 2018.

[5] https://wageindicator.org/salary/wages-in context. Living wages 2018, 2018.

[6] Wikipedia contributors. List of minimum wages by country — Wikipedia, the free encyclopedia. https://en.wikipedia.org/w/index.php?title=List_of_minimum_wages_by_country&oldid=969131742, 2020. [Online; accessed 24-July-2020].

[7] Rob Dellink, Jean Chateau, Elisa Lanzi, and Bertrand Magné. Long-term economic growth projections in the shared socioeconomic pathways. *Global Environmental Change*, 42:200–214, jan 2017.

[8] Keywan Riahi et al. The shared socioeconomic pathways and their energy, land use, and greenhouse gas emissions implications: an overview. *Global Environmental Change*, 42:153–168, 2017.
